# Bayesian Phylogenetic Inference of HIV Latent Lineage Ages Using Serial Sequences

**DOI:** 10.1101/2022.06.08.495297

**Authors:** Anna Nagel, Bruce Rannala

## Abstract

HIV evolves rapidly within individuals, allowing phylogenetic studies to infer the history of viral lineages on short time scales. Latent HIV sequences are an exception to this rapid evolution, as their transcriptional inactivity leads to negligible mutation rates in comparison to non-latent HIV lineages. Latent sequences are of keen interest as they provide insight into the formation, persistence, and decay of the latent reservoir. Different mutation rates in latent versus active HIV lineages generate potential information about the times at which sequences entered the latent reservoir. A Bayesian phylogenetic method is developed to infer integration times of latent HIV sequences. The method uses informative priors to incorporate biologically sensible bounds on inferences (such as requiring sequences to become latent before being sampled) that many existing methods lack. A new simulation method is also developed, based on widely-used epidemiological models of within-host viral dynamics, and applied to evaluate the new method, showing that point estimates and credible intervals are often more accurate by comparison with existing methods. Accurate estimates of latent integration dates are crucial in dating the formation of the latent reservoir relative to key events during HIV infection, such as the initiation of antiretroviral treatment. The method is applied to analyze publicly-available sequence data from 4 HIV patients, providing new insights regarding the temporal pattern of latent HIV integration events.

**Significance Statement:** Phylogenetic studies are increasingly being used to characterize within-host HIV evolution and the temporal dynamics of the HIV latent reservoir in particular, which is not targeted by current treatment methods and thus prevents a cure for HIV. Phylogenetic methods currently used to analyze HIV sequences suffer from conceptual and statistical problems that degrade their performance. A new Bayesian inference method to estimate the ages of latent sequences and a new simulation method based on within-host viral dynamics are developed. The new inference method outperforms existing methods, particularly in characterizing uncertainty. Understanding how the latent HIV reservoir changes overtime will allow researchers to better understand the nature of HIV infection and develop strategies for a cure.

---

A major obstacle to the development of a cure for HIV has been the presence of latently infected cells. HIV is a retrovirus that integrates its genome into the host cell genome. During latent infection, the integrated provirus is in a reversible state of transcriptional inactivity. Latently infected cells are not targeted by current treatment methods, namely antiretroviral therapy (ART). Consequently, treatment must be continued for life or the reactivation of latent cells will lead to a rapid rebound in viral load and disease progression (1). A detailed understanding of the dynamic processes of seeding, reseeding, and decay of the latent reservoir through the inference of latent integration dates for individual proviruses will allow researchers to have a better understanding of the nature of the reservoir as they work toward a cure for HIV.

HIV infects immune cells, specifically CD4+ cells, such as helper T cells and macrophages. Most infected cells die quickly (2, 3). In contrast, memory T cells have a long half-life of 4.4 years and can thus establish a latent reservoir for HIV (4). Memory T cells may be infected directly or an activated T cell may revert back to a quiescent state (5). Latently infected memory T cells can be activated by antigens, leading to the activation of the HIV provirus (6). Effective ART prevents infections of new host cells but does not prevent infected cells from producing virions. HIV can persist hidden in memory cells for decades, even with effective ART (4).

The latent reservoir is initially formed within days of infection and continues to be reseeded over time (7–9). However, the extent to which the composition of the reservoir changes over time is unclear. Some studies concluded that the latent reservoir that exists during ART is mostly seeded shortly before treatment initiation (10–12), while others have concluded that the reservoir is continuously seeded until treatment initiation (13). However, some of these results are difficult to interpret as a variety of mechanisms could account for these patterns. The timing of the formation of the latent reservoir is ultimately an empirical question that can be studied in multiple ways. In addition to further experimental work, reconstructing the ages of latent lineages can in principle be done by analyzing the patterns of variation observed among sampled sequences and applying phylogenetic methods designed to estimate sequence divergence times with serial sequence samples (11–16). The focus of this paper will be the development of new statistical and computational methods to accurately date the integration times of sampled latent sequences.

A variety of heuristic methods have been developed to estimate integration times using a combination of RNA sequences from serial sampled actively replicating sequences and RNA or DNA from putative latent sequences. All methods rely on a fixed estimate of the gene tree topology for the HIV sequences and some require branch lengths. Jones et al. developed a distance method that used linear regression (LR) to estimate the mutation rate from root-to-tip distances and sampling dates for non-latent sequences. This mutation rate is then used to estimate the latent integration dates (13). This method relies on a molecular clock, and is not used if the clock is rejected. Jones and Poon developed a related method, estimating mutation rate in the same way but estimated internal node ages and unknown tip ages using a maximum likelihood (ML) approach using a specified mutation rate (15, 16). To et al. developed a distance method using a least squares (LS) approach to estimate mutation rates and date internal nodes and tips with unknown ages (17). Their method requires the sequence length for estimating confidence intervals, but not the alignment. It was designed for extremely large phylogenies, but is applicable to HIV latency datasets as well. Abrahams et al. used multiple heuristic methods to date latent sequences. In one method, the distance from the closest sequence to the latent sequence, *d*, is determined, and the age of the latent sequence is assigned based on the sample time of the majority of sequences within 2*d* of the latent sequence (11). A similar method traverses the tree from the latent sequence toward the root of the tree until a node with 90% bootstrap support is found with at least one pre-treatment sequence. Then a latency time is assigned based on the most common sampling time of the pre-treatment sequences descendant from the well supported node (11). The two methods used by Abrahams et al. may be very sensitive to the number of sequences sampled and the sampling times. Simulation studies suggest that LS may out-perform all of these methods (15, 17). An alternative to these existing methods could be developed based on established parametric phylogenetic models that use tip dating for estimating and calibrating phylogenies of viral data, and are potentially more accurate (18, 19).

It has been difficult to evaluate the statistical performance of current methods for inferring integration times of latent HIV since existing simulation methods are biologically unrealistic. During the acute phase of infection, viral load grows exponentially shortly after infection, peaking within several weeks (20). Then the viral load falls one to two orders of magnitude before reaching a quasi-steady state. During this chronic phase of infection, the viral load remains relatively unchanged or rises only slowly until the onset of AIDS. In contrast, simulation methods that have been used to evaluate methods for dating integration events largely ignore the underlying population dynamics of HIV. Some assume a constant rate birth-death process while other use a compartmental model with logistic growth (13, 15). Epidemiologists use more complex models, typically ordinary differential equations (ODEs), to describe HIV viral dynamics (21–23). These models produce population trajectories that more closely match empirical observations, especially during acute infection, but the models have yet to be used in simulations to generate within-host HIV sequence data. The time period of acute infection is known to be important in establishing the latent reservoir (7), and this peak dynamic should be incorporated into simulation methods used to test inference methods aimed at estimating latency times.

We propose a Bayesian inference method to infer the latent integration date of HIV sequences. This is a full likelihood method, conditional on the phylogenetic tree topology. Additionally, we develop a simulation method based on existing viral dynamic models of HIV to test the performance of the inference method. The simulation model is parameterized using estimates from empirical datasets that produce realistic viral population dynamics (See SI section 4) (24).

## Model

A new program, HIVtree, was developed by modifying an existing program, MCMCtree, to infer latent integration dates (18). MCMCtree is a Bayesian phylogenetic inference program which estimates a time calibrated tree using viral sequences with serial samples given a fixed tree topology. It uses Markov chain Monte Carlo (MCMC) to estimate the model parameters. HIVtree incorporates additional parameters, the latent integration times, into the model. The program also estimates the originally defined parameters in MCMCtree, including substitution model parameters, substitution rate, and the internal node ages.

HIVtree assumes a priori that some sequences are known to be latent while others are not. Every sequence must also have a known sample date. In addition, every latent sequence has an unknown latent integration date. The youngest possible latent integration date is the sample time, and internal nodes cannot be latent. There is an optional bound on the oldest possible latent integration time, which could correspond to the oldest possible infection time. The model assumes that latent lineages have a mutation rate of zero, and all other lineages follow strict molecular clock. For calculating the likelihood, the latency time is treated as if it were the sample date for a non-latent lineage. This acts to reduce the tip age to be the time the sequence became latent (Fig. S4).

### Markov Chain Monte Carlo (MCMC)

HIVtree adds an additional step to the MCMC to estimate the latent times. In MCMCtree, proposals to non-root internal node ages are bounded above by the age of the parent node and below by the age of the oldest daughter node. A new time for each internal node is proposed within these bounds, the acceptance ratio is calculated, and the move is either accepted or rejected (18). In HIVtree, in addition to bounds on nodes, latent times are bounded above by the age of the parent node and below by the sample time. This ensures that the sequence becomes latent before it is sampled and that internal nodes cannot be latent. If the optional bound on latent integration times is used, the younger of the parent node age and the bound is used as the bound. Similar to MCMCtree, for each latent time, a move is proposed within these bounds, the acceptance ratio is calculated, and the move is either accepted or rejected (Fig. S4). Other than the difference in bounds, the proposal moves for the internal nodes and the latency times are identical. For the mixing step, the latency time is treated as equivalent to the sample date. The mixing step was not modified from MCMCtree (18).

### Prior Model

Two new root age priors were implemented in HIVtree. HIVtree and MCMCtree both require the user to specify the priors in backward time. The time of the last sample is considered to be time zero, and earlier times are positive. The programs also require a specification of a time unit transformation. For example, consider HIV data with the sample times specified in days. A time unit of 1000 days means that 0.365 is equivalent to a year in the prior specification. A shifted gamma prior, Γ(*α*, *β*), is implemented as the root age prior. The distribution is shifted by adding the first sample time to the distribution. This ensures there is no density after sequences are sampled. The gamma distribution parameters must also be chosen with the time unit transformation going backward in time. An option for a more informative prior is a uniform prior with narrow hard bounds (zero tail probability), U(*a*, *b*). There is no explicit prior on the internal nodes ages which is equivalent to a uniform prior on the possible node ages given the constraints from the sampling dates and the root age. Since the sampling prior is not explicit and the rank order of the nodes and the constraints jointly determine the prior, the MCMC must be run without data in order to recover the prior for the internal nodes, latency times, and root age. The distribution of the root age when the MCMC is run without data will not be equivalent to the user specified prior (Fig. S5). This effect is similar to constraints imposed by fossil calibrations (25). The mean root age will be older than the expectation of the prior distribution. The parameters of the gamma distribution can be modified to achieve a desired mean and variance for the root age. Using a uniform prior with a wide interval is discouraged due to this effect (an induced prior age of the root that is very old).

### Combining Inferences Across Genes

HIVtree only allows single locus inferences. However, the entire HIV genome is incorporated in the host cell genome at the same time, meaning different genes share the same latent integration times. Let *X* = {*x_i_*} be sequence data for *n* loci, where *x_i_* are sequence data at locus *i*. Let *T* be a latency time that is shared across loci. The remaining parameters of the gene tree may be different due to recombination. The posterior density of *T* is

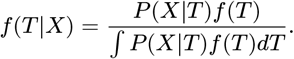

If we ignore the correlation between gene trees due to limited recombination and treat the loci as independent the posterior density can be written as

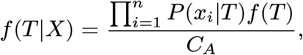

where *C_A_* is the marginal probability of the data (which is a constant),

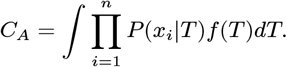

We want to calculate the posterior probability of *T* for each locus separately using MCMC and subsequently combine them to obtain a posterior density for all the loci. To do this we formulate the above equation as a product of the marginal posterior of *T* for each locus,

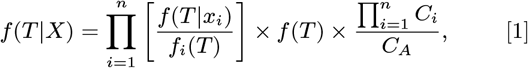

where *f_i_*(*T*) is the prior on *T* for the *i*th locus and *f*(*T*) is the desired prior for the combined posterior. The last term is a proportionality constant that insures the posterior density integrates to 1. A simple example illustrating this general approach to combine posteriors using a normal distribution is provided in SI section 8.

In our analyses, *n* independent MCMC analyses are run (with and without using the likelihood) and kernel density estimation is used to estimate *P*(*T*|*X_i_*) and *f_i_*(*T*), respectively, for *i* = 1,…, *n*. The estimated kernel functions are then used to evaluate equation 1 up to an unspecified proportionality constant (see supplemental material). Simulations were used to evaluate the performance of this approach to combine posteriors.

## Results

### Simulation Analysis

Here we compare the statistical performance of HIVtree and several other existing methods when analyzing simulated datasets with known latency times.

#### Comparisons on a Fixed Tree Topology

HIVtree was compared with three existing methods, least squares dating (LS) (17), linear regression (LR) (13), and pseudo maximum likelihood (ML) (16) using simulated datasets. The effect of variation among the independently simulated sequences on point estimates of latent tip ages can be seen by comparing the estimates for a given latent tip in a fixed tree. Even with *C1V2*, the most informative gene simulated, there is considerable variation in the estimated latency time for a given latent tip (Fig. 1). The variation is even larger for the other genes (Fig. S6). The estimated times for a single latent tip sometimes differs from the true value by a decade or more for both the LR and ML methods. The LS method has fewer extreme estimates, which are prevented by bounds on the integration times. LS allows for upper and lower bounds for each individual latent sequence while ML has the same upper bound on all latent sequences, which is the last sample time. The LR has no bounds on the inferred integration time, potentially allowing the latent sequences to be formed either after the sequence was sampled or before an individual was infected. Both outcomes are logically impossible.

**Fig. 1.**
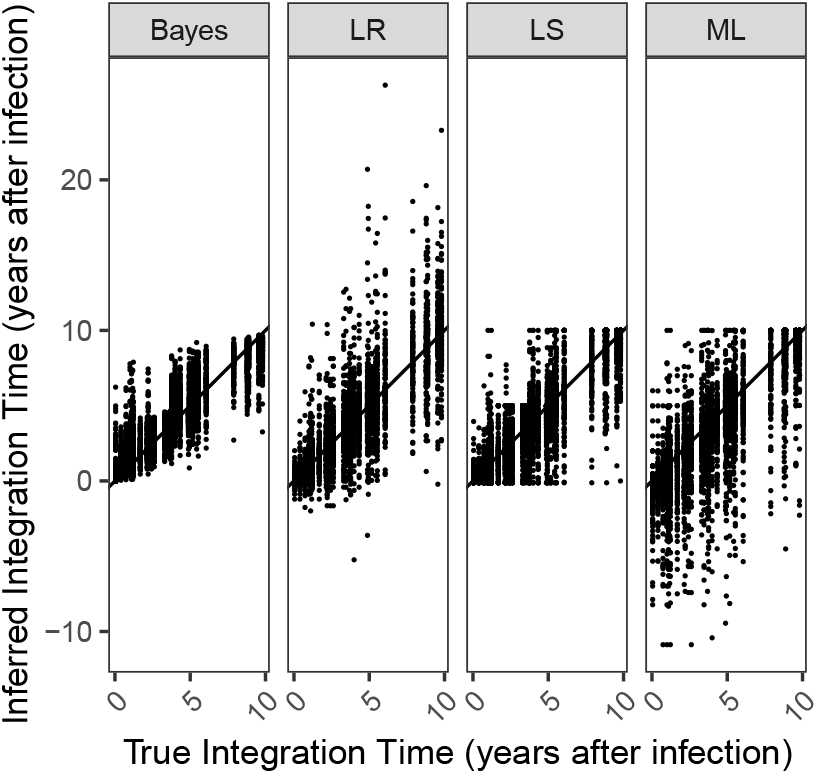
For all 30 alignments simulated for *C1V2* on a fixed tree, the inferred integration dates are shown for each method. If the methods performed perfectly, all points would fall on the line, which is has an intercept of 0 and slope of 1. The units are years after infection.

#### Inferences Across Genes

The posterior distribution for each latent time is inferred separately for each gene when using HIVtree. When the marginal densities are combined across the genes, the posterior densities become narrower and closer to the true value (Fig. 2). The other methods do not allow such information sharing.

**Fig. 2.**
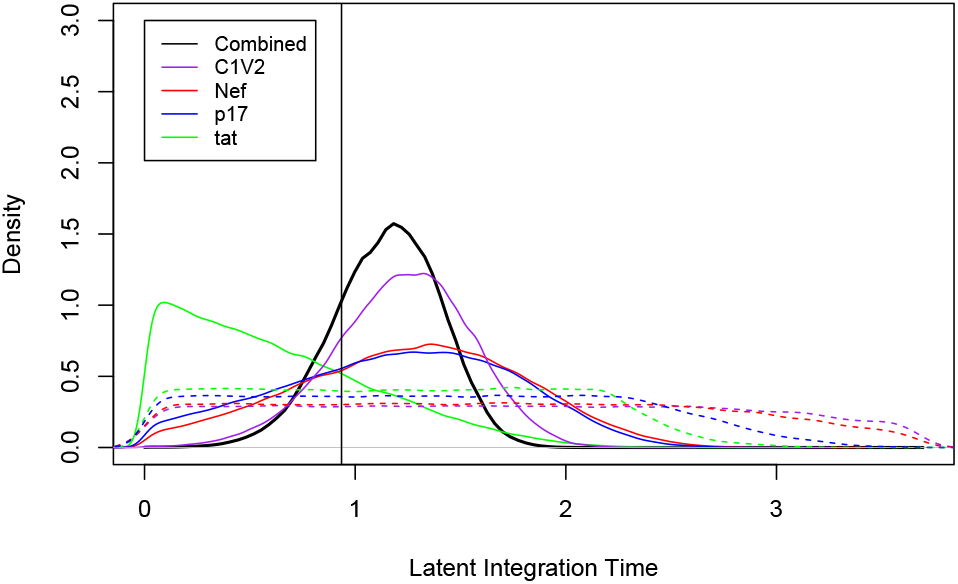
Joint posterior density for a single latency time across all genes. Each solid colored line shows the marginal posterior density for a single latency time for different genes. The dashed colored lines show the marginal prior densities, which result from running the MCMC without data. The solid black line shows the estimate with the genes combined. The vertical line is the true latent integration time. The MCMC was run for 500,000 iterations, sampling every other iteration. This results in smoother curves than the shorter MCMCs run used in the larger analysis of simulated data, but results are very similar.

#### Summary of Method Performance

Mean square error (MSE) is a useful measure of method performance that includes both bias and variance and is directly comparable across methods. MSE is lowest for *C1V2* and highest for *tat* for all analyses (Fig. 3a). All of the methods are the least biased for *C1V2* and the most biased for tat (Fig. 3b). The average bias for the ML and LS methods are more negative for the shorter, slower evolving genes, while the Bayesian and LR method have a positive bias on average.

**Fig. 3.**
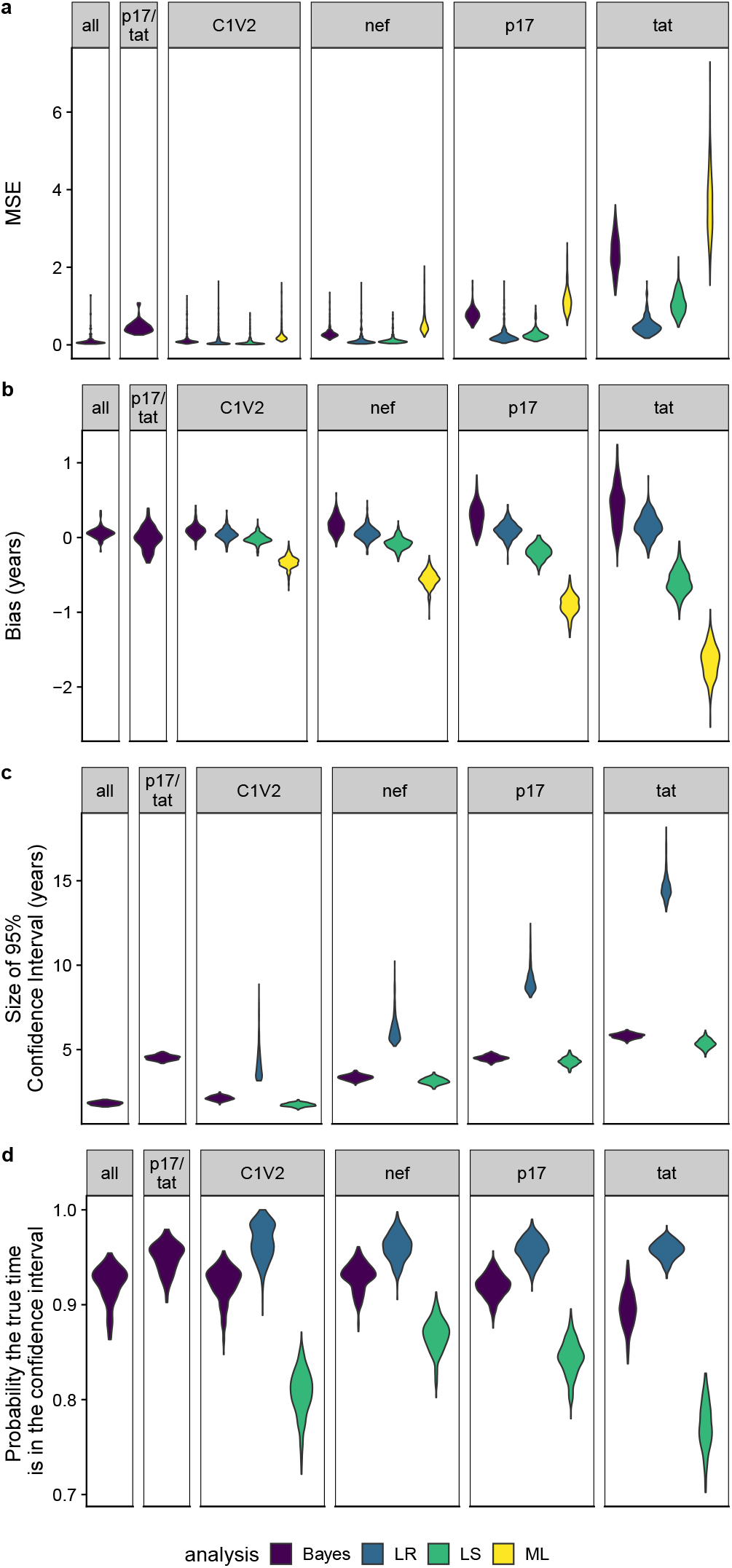
For each of fixed tree topologies, the mean square error (MSE), bias, and size of the 95% confidence/credibility interval was averaged across all 900 latent times for each gene analysis combination. Each violin plot is made using 300 data points, corresponding to the average from each of the 300 fixed tree toplogies. For the Bayesian combined analysis of either all of the genes or only p17/tat, only a third of the fixed tree toplogies were analyzed.

In the simulation analysis, the probability that the true value falls in the 95% confidence interval (or 95% highest posterior density for Bayesian analysis) is considered (Fig. 3d). The Bayesian method has comparable coverage probabilities for *C1V2* and *nef* of 92% and 93%, respectively, with the lowest coverage probability for *tat* (90%). The average size of the 95% credible set for the longest and shortest sequences, *C1V2* and *tat*, is 2.1 years and 5.8 years, respectively. The LR has the highest coverage, with a coverage probability of 97% for *C1V2* and 96% for *tat*. However, LR has very large confidence intervals (Fig. 3c). The mean size of the 95% confidence interval is 4 years and 15 years for *C1V2* and *tat*, respectively. In contrast, the LS method shows lower coverage probabilities but smaller confidence intervals. The LS method has its highest average coverage probability for *nef* (87%), but drops to 77% for *tat* (Fig. 3d). For the longest gene, *C1V2*, the average coverage probability is only 81%. This is likely due to the much smaller confidence interval size. The size of the 95% confidence interval is much larger for the LR method than either the LS or Bayesian methods (Fig. 3c). The LS and Bayesian methods have similar size confidence intervals, but the Bayesian method is more likely to contain the true value in the 95% confidence interval (has higher average coverage probability). The ML method has the largest MSE and bias on average for all regions and does not provide confidence intervals.

When the inferences are combined across all four genes, the average size 95% credible set is 110 days smaller on average. The average probability the true integration time is in the 95% credible set is very similar to the results for the longest gene. When the two shortest genes, *p17* and *tat*, are combined, the average size of the 95% credible set is very similar to *p17* alone, but the probability the true value is in the 95% credible set increases from 92% with *p17* alone to 95% in the combined analysis (Fig. 3c,d).

### Empirical Analysis

We applied each of the four methods to HIV data sets from two studies of serial sampled HIV sequences. The first data set (Jones et al.) is comprised of *nef* sequences for two patients (13). For each patient, plasma HIV RNA was sequenced multiple times over a period of almost a decade either pre-treatment or during incompletely suppressive dual ART. After the initiation of combination ART (cART), samples from the putative reservoir were taken from at least two time points. Samples consisted of HIV RNA sequences sampled during viral blips and proviral DNA collected from whole blood and peripheral blood mononuclear cells (PBMC). The second data set (Abrahams et al.) has three regions of *env* for both the patients analyzed (217 and 257) and *gag* and *nef* sequences for one patient (257) (11). For both patients, virus was sequenced from the plasma multiple times over several years prior to ART initiation. After ART initiation, viral RNA was isolated from the supernatant of quantitative viral outgrowth assays.

The inferred latent integration times for the patients in the Jones et al. dataset obtained using HIVtree span over a decade (Fig. 4), similar to estimates obtained using other methods (Fig. S7). However, ML and LR infer integration times that occur after the sampling time in some cases (Fig. S9). For the Abrahams et al. dataset, the point estimates, especially for the early sample times (11.1 for patient 1 and 17.9 for patient 2), tend to be concentrated near the time of ART initiation. The combined point estimates for the latency times inferred using HIVtree appear loosely clustered around the time ART began for patient 257, with narrower credible sets than the analyses on individual genes (Fig. 5). These patterns for patient 217 are less clear, possible due to fewer genomic regions and fewer latent sequences (Fig. S8). Sometimes LS gives very large confidence intervals, covering the entire area between the bounds for a sequence (Fig. S10, S13), while in other cases the confidence intervals are smaller than LR.

**Fig. 4.**
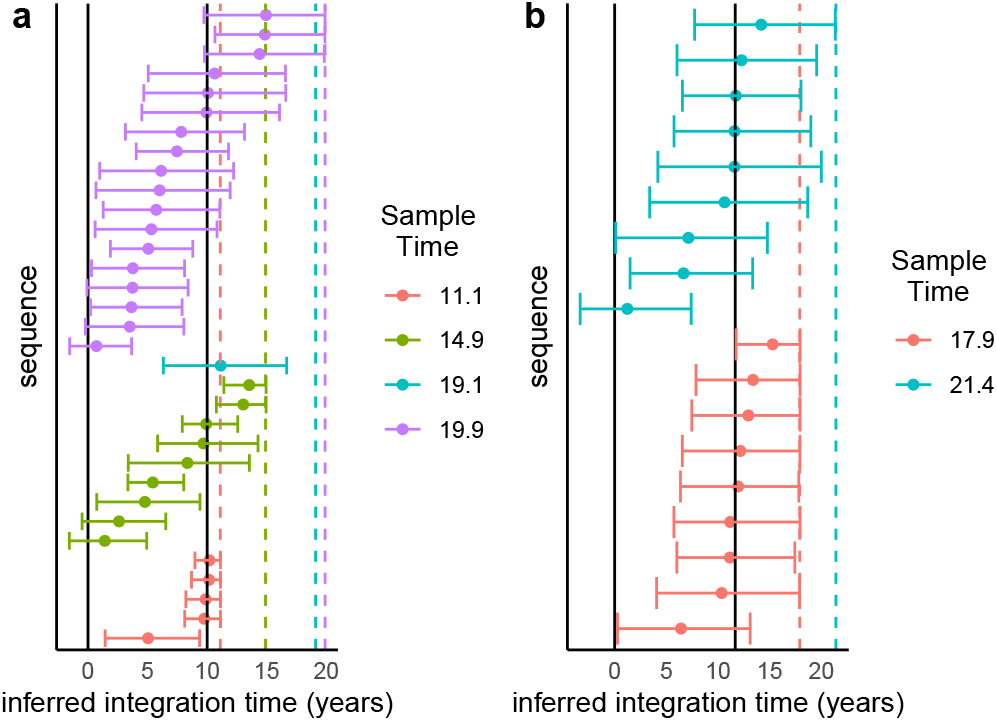
Panels (a) and (b) show the inferred latent integration times, in units of years after diagnosis, for patients 1 and 2, respectively, inferred using HIVtree to analyse sequence data for the *nef* gene locus. A dot indicates the posterior mean and bars represent the 95% credible interval. The solid vertical lines indicate the positive test date (left) and time of cART initiation (right) for each patient. The colored dashed vertical lines indicate the sample times.

**Fig. 5.**
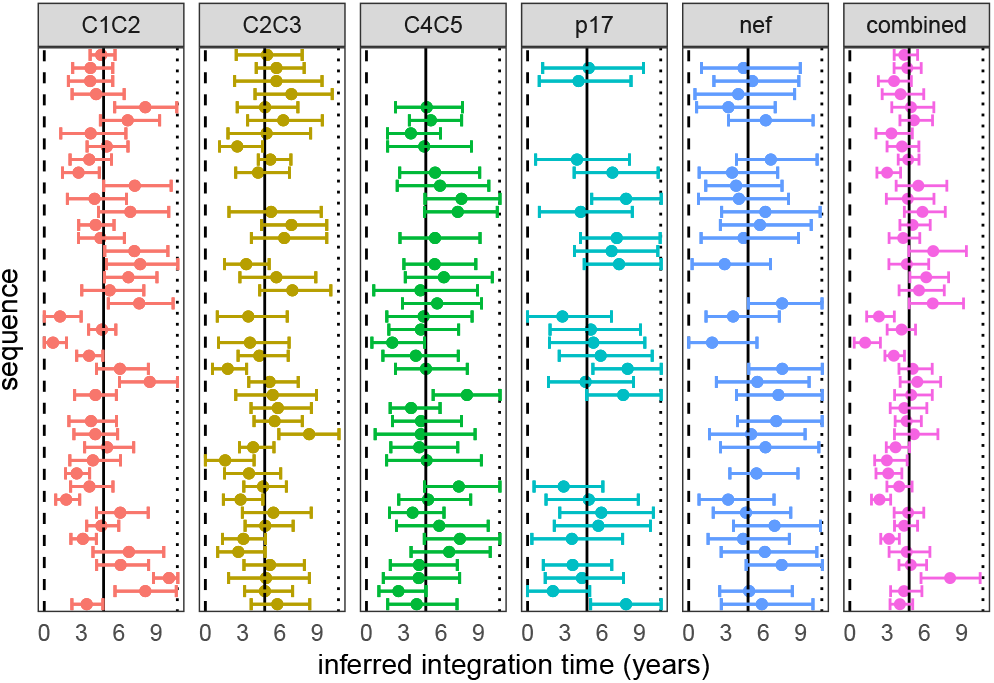
The five panels to the left each show the integration times inferred using HIVtree for a single gene locus. The panel to the right shows the inferred integration times when posterior distributions for the five loci are combined. A dot indicates the posterior mean and bars represent the 95% credible interval, in units of years after diagnosis. The results are from patient 257 (11). 10 non-latent sequence were used as each available timepoint and sites with more than 75% gaps were removed from the alignment prior to analysis, as described in SI section 10. The dashed line shows the infection time, the solid line shows the start of ART, and the dotted line shows the sample time.

## Discussion

Here, we have described both a phylogenetic method to infer latent integration times and a new method to simulate sequence data based on within-host viral dynamics. HIVtree performs better than existing methods by a variety of metrics. The method has smaller confidence intervals on average than alternative methods, while still containing the true value, resulting in more precise interval estimates of the integration dates. Moreover, the MSE is comparable to the best alternative method when the data are informative.

HIVtree has several improvements over existing methods. It allows for biologically relevant bounds on latent integration times, such as requiring the latent times be older than the sample times with an option to bound the integration times at the time of infection. Among the alternative methods, only the LS method allows for such bounds. Bayesian inference also provides a sensible way to combine estimates across genes, while allowing for potentially different gene tree topologies. This results in more precise estimates, especially when the sequences available are short. There is currently no alternative to the HIVtree method for jointly inferring latency times using multiple loci, nor is there a clear way to do so. Lastly, Bayesian methods have the advantage of well known statistical properties, such as statistical efficiency and consistency. By treating an alignment as data, HIVtree allows for full use of the available sequence data in the inference, whereas the other methods only use an inferred phylogenetic tree which may not be a sufficient statistic.

There are several avenues for improvement of HIVtree. In the current paper, to use data from multiple loci in HIVtree the marginal distributions for the latent integration times were combined. A more formal method to combine data across loci would be to jointly analyze the loci in a single model, allowing the MCMC to integrate over the node ages in each of gene trees separately while constraining the latent integration times to be the same for sequences derived from an individual infected cell. This would be most sensible to implement in a program that accommodates multilocus data, such as bpp (26), rather than the parent program of HIVtree, mcmctree.

Further, despite desiring a diffuse prior on the node ages and latent times, the prior model in HIVtree seems to be too informative in some cases. The rank order of the nodes and the serial sampling cause average the root age of the phylogeny in the prior to be older than the user input prior. If the root age is constrained, such as by using a uniform prior, the latent times are pushed closer to time present, which introduces a bias to the latent inferences (unpublished preliminary analysis). This means that constraining the root age to be close to the true age can produce worse estimates of the latent times. Similar effects driven by constraints among node ages have been previously noted for fossil calibrations and serially sampled data (18, 27). However, the effects appear to be more pronounced when the root ages are close to the the serially sampled sequences, as can result from within-host viral data. While there may be quite informative outside knowledge on the age of the root for HIV, such as the time of infection, we currently caution against forcing the root age to match the infection time when using HIVtree because this may induce bias in estimates of latent virus integration times.

The difference between the user input prior distribution on the root age and the prior observed when running the MCMC without data appears to be larger with the empirical data than with the simulated datasets. While the exact cause of this discrepancy is unknown, it may be related to the ladderlike tree topologies of the empirical data or the sampling times of the sequences. A different prior may improve some of these limitations. One option would be a serial sample coalescent prior with changing populations sizes (28, 29). This would also be more sensible to implement in a program which includes coalescent models, such as bpp. Such a prior could also allow for the incorporation of information on viral population sizes (such as from well described viral dynamic models) and knowledge of the time of infection.

The viral dynamic simulation method developed in this paper is based on well-studied models of HIV population dynamics within hosts. This is likely to be more realistic than traditional methods used to simulate phylogenies, such as constant rate birth-death processes, and it follows standard epidemiology approaches for studying viral dynamics. However, this model does not incorporate selection, which is known to be important in HIV evolution. The method produces trees that are more star-like, with short internal branches, than those typically inferred in empirical studies of HIV sequences. Future work should focus on modeling selection, as well as other aspects of HIV biology, such as clonal proliferation of latently infected immune cells, to develop simulators and priors for inference that more accurately model HIV biology and produce trees that more closely match the empirical observations.

## Materials and Methods

Here we provide a brief description of the materials and methods used in this paper, which are described fully in the SI Appendix.

### Simulation of Phylogeny

A stochastic simulation based on existing ODEs was developed to simulate tree topologies of sampled latent and active HIV sequences. In the ODE, the sizes of five populations of cells and viruses are tracked, including uninfected CD4+ target cells, productively (actively) infected CD4+ target cells, virions, replication-competent latent cells, and replication-incompetent latent cells (see SI section 1). The stochastic model is formulated as a continuous-time Markov chain with instantaneous rates as described in the deterministic model (see SI section 2). The process is modeled as a jump chain. A user specified number of virions and latent cells are sampled at any number of user specified times.

A C program was written to to simulate under the stochastic model. In addition to simulating population sizes, it tracks the parent-daughter relationships of all infected cells and viruses in a binary tree (see SI section 3). The amount of time latent in each branch is also tracked. The stochastic and deterministic models are in good agreement when population sizes are large, as expected (Fig. S3). The total number of tips in the tree varied over time. The maximum number of tips in a tree was on the order of 10^8^ (Fig. S3).

### Simulation of Sequence Data

A separate C program was written to simulate DNA sequences given a sampled tree with branch lengths and a latent history. Sequences are simulated in the typical manner, assuming independent substitutions among sites, starting at the root of the tree and simulating forward in time toward the tips of the tree. The simulator accommodates models as general as the GTR+Γ substitution model (30, 31). No substitutions can occur while a lineage is latent. The program allows an outgroup with a node age of zero to be simulated. The sequence at the root is specified by a FASTA format input file (from an existing HIV sequence, for example).

### Sampling and simulation parameters

100 trees were simulated using the stochastic simulator. 50 viruses and 10 latent cells are sampled every year for 10 years. On the tenth year, an extra 50 latent cells are sampled. For each of these 100 phylogenies, 30 alignments for each of four genomic regions were generated with the DNA simulator using an outgroup. To determine the DNA substitution parameters, within-host longitudinal samples from published data sets for four regions (*tat*, *p17*, *nef*, *C1V2*) were analyzed with MCMCtree (see SI section 6). The estimated substitution rate and length varied among the simulated regions, with *C1V2* having the highest substitution rate (*μ* = 3.56 × 10^−5^ per base per day) and the most sites (*n* = 825) and *nef* having the next highest substitution rate (*μ* = 1.34 × 10^−5^ per base per day) and number of sites (*n* = 618). *p17* has a slightly lower substitution rate than *tat* (*μ* = 8.9 × 10^−6^ per base per day versus *μ* = 9.9 × 10^−6^ per base per day), but more sites (*n* = 391 versus *n* = 132)(Table S2). For each phylogeny and alignment, the sequences and phylogenies were then subsampled three times to generate three trees and three corresponding alignments. Specifically, 10 viruses were subsampled every year for 10 years. 10 latent cells were subsampled after 5 years of infection and 20 were subsampled after 10 years of infection. In total, 300 tree topologies were simulated, each with 30 latent and 100 non-latent randomly sampled sequences. This led to a total of 300 topologies × 30 alignments × 4 regions = 36, 000 simulated datasets.

### Maximum Likelihood Tree Inference and Rooting

To analyze the simulated datasets a rooted tree topology was first inferred for use by HIVtree and other heuristic programs. Maximum likelihood trees were inferred with raxml-ng using an HKY+Γ model and outgroup rooted (32, 33). 25 parsimony and 25 random starting trees were used for the tree search. The outgroup was removed from the inferred tree. Both the LS and Bayesian methods use the outgroup rooted tree. For the ML method, the tree was re-rooted using root to tip regression available in the R package ape prior to analysis (19, 34). The LR method re-roots the tree using root to tip regression as part of the analysis. For LS, the sampling time was used as an upper bound for the latent lineages and the lower bound was 45 days prior to infection, while the active lineages were constrained to their sampling time. The ML and LR methods do not include additional constraints.

### Bayesian inference

For HIVtree analyses of simulated data, an HKY+Γ model was used with 5 rate categories and the prior *κ* ~ G(8, 1) (32). The prior for among site rate variation was *α* ~ G(4, 8). A time unit of 1000 was used with a substitution rate prior of G(2, 200), meaning the mean was 10^−5^ per base per day. The root age prior was Gamma(36.5, 100). The latent times were bounded at 3.695, which is equivalent to 45 days prior to infection. Two MCMCs were run for each analysis to check for convergence. MCMC lengths and conditions for convergence are described in the SI Appendix (see SI section 7).

### Combining Posterior Estimates from HIVtree

For combining results in Bayesian analyses of the simulated and empirical datasets, the function kdensity in the kdensity R package was used for kernel density estimation of the posterior distribution and the prior distribution of each latent time (35). The posteriors and priors for each gene were multiplied according to equation 1. The resulting function was normalized by finding the proportionality constant using the integrate function. For the simulated datasets, the integral bounds were set to the bounds on the latent time in HIVtree, which was the sample time and 45 days prior to infection. The 0.025 and 0.975 quantiles were found using the invFunc function in the R package GoFKernel (36). The mean for the joint posterior was found using the integrate function. For the simulated datasets, this analysis was conducted on only a third of the trees from the main simulation analysis due to the highly demanding computations involved. For a small subset of simulated data, numerical issues prevented estimation of a combined latent integration time. (see SI section 8b).

### Existing Methods

The LR method was run using scripts available at: https://github.com/cfe-lab/phylodating The ML method used scripts available at: https://github.com/brj1/node.dating/releases/tag/v1.2 The driver script provided by Jones et al. is available at: https://github.com/nage0178/HIVtreeAnalysis The LS method was obtained from: https://github.com/tothuhien/lsd-0.3beta/releases/tag/v0.3.3

### Empirical Analysis

Data sets published from (11, 13) required curation prior to analysis. Due the large number of sequences in the the Abrahams et al. data set, sequences were subsampled, and alignments were edited due to gaps (see SI section 10). For all empirical data sets, raxml-ng was run using an HKY+Γ model (33). 25 parsimony and 25 random starting trees were used for the tree search. Trees were rooted using root to tip regression using the rtt function in the ape package available in the R package ape prior to analysis (19, 34). Each of the four methods were run on all datasets.

For the Jones et al. dataset, HIVtree was run with a root age prior of G(8,60) for patient 1 and G(15,50) for patient 2. These priors were chosen to have an induced prior when running without data with a variance of several years and a mean several years prior to diagnosis. Latent integration times were bounded 10 years prior to diagnosis, as a very conservative oldest possible bound. In the HIVtree analysis, an HKY+Γ model was used with 5 rate categories with the prior *κ* ~ G(8, 1). The prior for among site rate variation was *α* ~ G(4, 8). A time unit of 1000 was used with a substitution rate prior of G(5, 1000), meaning the mean was 5 × 10^−6^ per base per day. For the LS analysis, latent integration times had the same bounds of 10 years prior to diagnosis and the sample times.

For the Abrahams et al. dataset, the LS and HIVtree analyses bounded the latent times at the infection times and the sample times. In the HIVtree analysis, an HKY+Γ model was used with 5 rate categories with the prior *κ* ~ G(8, 1). The prior for among site rate variation was *α* ~ G(4, 8). A time unit of 1000 was used with a substitution rate prior of G(2, 200), meaning the mean was 10^−5^ per base per day. The root age prior was G(0.25, 110) for all datasets. This prior was chosen to have a relatively wide variance on the root age with a mean slightly before the infection time as well as a large variance on the latent integration times. As described in the Prior Model section, the root ages are older than the given prior when run without data, and they are also different for each dataset. When running the MCMC under the prior, small changes to the prior appeared to cause little change to the posterior distribution of the latent integration times. A full description of the MCMC convergence criteria is provided in SI sections 9 and 10 for the Jones et al. and Abrahams et al. datasets, respectively. The Jones et al. dataset only sampled one gene, so estimates from multiple genes could not be combined. The estimates from multiple genes for the Abrahams et al. dataset were only combined for the tree with 10 non-latent sequences per sampling time and sites with gaps in over 75% of the sequences were removed from the alignment.

### Program availability

The gene tree and the DNA simulation software packages are available at: https://github.com/nage0178/HIVtreeSimulations

The HIVtree software package is available at: https://github.com/nage0178/HIVtree

Scripts to produce the results in this paper are available at: https://github.com/nage0178/HIVtreeAnalysis

## ACKNOWLEDGMENTS

A.N. was supported by the National Science Foundation Graduate Research Fellowship Program under Grant No.2036201. This research was supported by National Institutes of Health Grant GM123306 to B.R.

## Supplementary Information

### Supporting Information Text

#### 1. Deterministic Model

Here we describe the deterministic model of HIV population dynamics that will serve as the large-population analog of our stochastic model (see below). Let *T*(*t*) be the number of uninfected target cells at time *t*. Let *T*\*(*t*) be the number of productively infected cells at time *t*. Let *L*(*t*) be the number of latently infected, replication-incompetent cells at time *t*. Let *L*\*(*t*) be the number of latently infected, replication-competent cells at time *t*. Let *V*(*t*) be the number of virions at time *t* (S1). Actively infected target cells that are replication-incompetent are not modeled. Define λ to be the rate at which uninfected target cells are produced and *d* to be the per cell rate at which they die. Let *δ* be the per cell rate at which actively infected cells die. Latent replication-competent cells and replication-incompetent cells die at constant per cell rates of *σ* and *τ*, respectively. Let ~ be the proportion of newly infected cells that are replication-incompetent. Let *η* be the proportion of newly infected cells that are latently infected and (1 – *η*) be the proportion of newly infected cells that are actively infected. Let *κ* be the rate constant for target cells becoming infected cells. Productively infected cells must be replication-competent and are produced at a rate equal to product of the rate constant *κ*, the number of virions, the number of uninfected cells, the proportion of cells that are replication-competent, and the proportion of cells that are actively infected. The rate of production of latent replication-competent cells is calculated similarly, except that the proportion of cells that are latently infected is used rather than the actively infected population. For replication-incompetent latent cells, the rate of production is equal to the product of the rate constant *κ*, the number of virions, the number of uninfected cells, the proportion of cells that are replication-incompetent, and the proportion of cells that are latently infected. When an infected cell is produced, an uninfected cell is lost, since the uninfected cell becomes the infected cell. This is true for actively infected cells and both types of latently infected cells.

Latent replication-competent cells can reactivate and become actively infected cells. This occurs at a constant per cell rate of *α*. HIV virions, *V*, are produced at a rate proportional to the concentration of actively infected cells, with rate constant *π*. The virions are cleared at a constant per virion rate of *c*. This model gives the following set of equations:

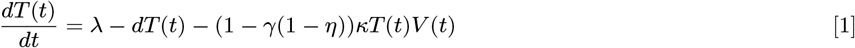

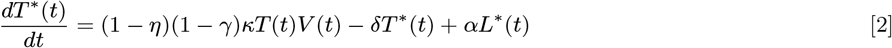

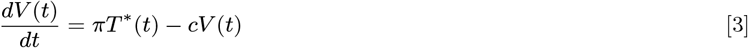

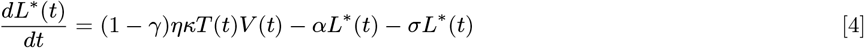

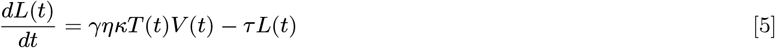

The solutions to these equation are obtained by numerical analysis using the function ode in the R package deSolve (1).

#### 2. Stochastic model

Viral dynamics were modeled using a continuous-time Markov chain with instantaneous rates as previously described in the deterministic model. For example, let *A* be the event that a birth of an uninfected cell occurs in the time interval Δ*t*. Then,

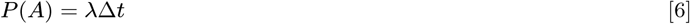

The process is modeled as a jump chain. Only one event can occur in a small interval Δ*t*, and the number of viruses, or of any cell type, can only change by one in that interval. The waiting time between birth events of uninfected cells is exponentially distributed with mean waiting time 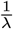. The instantaneous rates and waiting time between other events are determined similarly. The total rate of events, *R*(*t*), is given by the sum of the rates of all possible events.

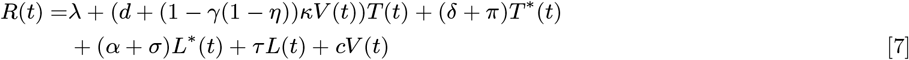

The waiting time between any event is exponentially distributed with mean 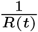. Given that an event occurs, the probability the event was a birth of an uninfected cell, for example, is given by the ratio of the rate of birth events of uninfected cells and the total rate of events, 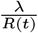. The probabilities of other events are determined similarly.

#### 3. Simulation of tree topologies

The stochastic model was implemented as a C program. In the program, the parent daughter relationship of all of the viruses in a tree structure is tracked. The cell or virus type (e.g. T*, V, L, or L*) is also tracked. The simulation is initialized with a single actively infected cell. Each time a virus is born, an actively infected cell is randomly selected to branch into two daughter lineages. One lineage is an actively infected cell and the other an active virus. Each time a virus or cell dies, an existing virus or cell of that type is randomly removed from the tree. When a virus latently infects a cell, a virus is randomly chosen to branch into an infected cell and a virus. This is designed to follow the conventional ODE models, even though a single virus cannot infect multiple cells in real systems. This is likely inconsequential, since the waiting time for a virus to die is short, and thus the probability a virus infects multiple cells is very small. Replication-competent latent viruses may be reactivated, meaning they become actively infected cells. Extinction is considered to be analogous to a failure to establish infection. In this case, the simulation is restarted. At pre-specified times, a pre-specified number of active viruses and latently infected cells are sampled. Replication-competent and incompetent cells are not distinguished during sampling. Sampling is equivalent to a death event for all sampled lineages.

#### 4. Parameter Values

Parameter values were determined using empirical estimates. Since many of the parameters are not independent and choosing parameters independently can lead to unrealistic patterns of viral load change over time, parameters obtained from a single patient and study were used for as many of the parameters as possible (2). The remaining parameters are taken from the literature (S1). *η* is fixed such that there are 1.4 × 10^6^ replication competent latent cells in 5L of blood at equilibrium (3). The initial concentration of uninfected target cells is assumed to be 10 cell/μL (2). Initially there is a single actively infected cell. All other cell and virus populations have size zero.

In principle, the simulation method described above would allow the entire viral population within a host to be simulated. However, this is not computationally tractable due to the simulation time and memory usage. ODEs of viral dynamics in HIV typically describe the changes in concentrations of cells and viruses per mL of blood. If properties of the viral genealogies become independent of the simulation size as the simulation size increases, it may be reasonable to use a simulation volume much smaller than the total blood volume in an adult. To determine whether this was the case, the impact of simulation size was examined by simulating genealogies generated with different blood volumes while keeping the number sampled sequences constant. Tree length increases and then plateaus as the simulation size increases. Other tree metrics, including root age and total time spent in latency, also showed no trend with volume (S2). Thus, 100 mL was used as the simulation volume.

#### 5. Agreement between the deterministic and stochastic models

For large population sizes, the stochastic model and the deterministic (ODE) model are expected to produce similar results for the population size as a function of time given the parameters and initial values are such that the population does not go extinct in the stochastic simulation. This is because we have designed the stochastic simulator to have an expected population size equal to the predicted population size for the deterministic model at any point in time and the relative variance of the stochastic model decreases with increasing population size. Populations sizes are in good agreement when there is no extinction (S3). Cases of extinction are common, but are not considered further.

#### 6. Estimation of DNA substitution model parameters

To select DNA substitution model parameters to use in the simulations, parameters were inferred from empirical datasets for four genomic regions using MCMCtree (4). Alignments for *nef*, *tat*, *C1V2*, and *p17* were taken from a studies on longitudinal Cytotoxic T-lymphocyte (CTL) responses from the LANL HIV special interest alignments (5–7). This patient (code PIC1362) was infected in 1998, was a homosexual male, and participated in a study at University of Washington Primary Infection Clinic. The patient had sequences samples taken at 18 time points and was untreated at the time of the study.

To root the tree, sequences from four patients were selected using the [LANL database](https://www.hiv.lanl.gov/components/sequence/HIV/search/search.html) to use as outgroups (GenBank accession numbers: AY331284, AY331289, AB078005, JN024426). The best outgroup is not always clear in phylogenetic studies. Multiple outgroups were used to compare of the effect of rooting on substitution rate estimates. All four of these patients were infected within 2 years of PIC1362, were likely infected on the west coast of the United States, has sexual transmission as a risk factor, were untreated at the time of sampling, and had all four genomic regions were available. The outgroup sequences were combined with the existing alignments using the [SynchAlign](https://www.hiv.lanl.gov/content/sequence/SYNCH_ALIGNS/SynchAligns.html) tool on the LANL HIV database. This resulted in 16 alignments, one for each gene outgroup pair. Then, sites with more than 75% gaps were removed from the sequences using a custom R script. This was done to remove problematic regions of the alignments, particularly in *C1V2*.

To obtain parameter estimates, maximum likelihood trees were inferred with RAxML-ng (8) under an HKY+Γ model (9, 10) and outgroup rooted. The outgroups were removed from each of the alignments and the maximum likelihood trees. MCMCtree was used to infer the substitution model parameters and substitution rate for each gene with each outgroup rooting (4). An HKY+Γ model with 15 rate categories was used. The prior for *κ* in the HKY model was G(8, 1). The prior for among site rate variation was *α* ~ G(1, 1). A time unit of 1000 was used with a rate prior of G(2, 200), or 10^−6^ substitutions per base per day. A birth-death-sequential-sampling model was used with parameters *λ* = 2, *μ* =1, *ρ* = 0, and *ψ* = 1.8 (11). A root age prior was U(1, 10), meaning the root age was 1000 to 10000 days prior to the last sample time, with 0.01 tail probabilities (12).

5 replicates of MCMCtree were run for each gene outgroup pair. Each MCMC was run with a burnin of 1000, sample frequency of 2, and 10000 samples. The estimates from each of the 5 replicate MCMCtree runs were similar in all cases, indicating the MCMC converged. The point estimate of the substitution rate and the 95% HPD interval bounds for the substitution rate were averaged over the 5 replicates. In most cases, each outgroup produced similar mutation rate estimates for a given gene. The outgroup rooting with the smallest 95% HPD interval of the substitution rate divided by substitution rate was used to provide parameters for DNA simulation. However, for *nef*, outgroup 1006 had a much different rooting than the other outgroups. CS2 and PIC55751 had the same root location. Of those two, the one with the smaller 95% HPD interval of the substitution rate divided by substitution rate was used. This resulted in JN024426 being selected as the outgroup for all genes. The first replicate MCMC run of MCMCtree with JN024426 as the outgroup rooting was used for parameters estimates for each gene. This included the estimates of *α*, *κ*, *μ*, and the stationary frequencies (S2).

The HXB2 sequence was used at the root sequence for the simulation of each region (S2). However, no bases were removed inside the sequence, as done in the original alignment in regions with over 75% gaps. An HKY model was used for the simulation since the parameters inferences were made with an HKY model MCMCTree.

#### 7. MCMC settings for Simulation Analysis

For each of the simulated datasets, HIVtree was run with two seeds. The MCMC was sampled every other iteration for 30,000 samples with a burn in of 2,500. Thus a total of 30000 × 2 + 2500 = 62, 500 iterations were run. The internal node ages of the two replicate MCMCs were compared for each analysis. If the mean age difference between the two replicate MCMCs was more than 10 days for more than 10 internal nodes, 20 days for more than 5 internal nodes, or 100 for any internal nodes, the MCMCs are considered to not have converged. A total of 347 pairs of MCMCs did not converge out of 36,000 pairs run. For each pair of MCMCs that did not converge, another 2 MCMCs were run with different seeds with 60,000 samples. Of those, 18 pairs of MCMCs did not converge. Those MCMCs were rerun again with different seeds, a burnin of 10000 iterations, and were run for 240,000 iterations, sampling every other iteration. All of these runs met the above convergence criteria except one. This was a simulated *nef* dataset and was removed from all analyses.

#### 8. Combining Posteriors

##### A. Example: Sample from a Bivariate Normal PDF

Suppose that we have samples *Y* = *y*_1_,…, *y_a_* and *X* = *x*_1_,…, *y_b_* from a bivariate normal density with means *μ_y_* = *μ_x_* = *μ*, variances 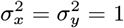 and correlation parameter *ρ*. Our goal will be to generate the posterior density of *μ* by combining posterior densities for *x* and *y*. We will treat the variables *Y* and *X* as independent in our inference procedure, though in reality *ρ* may be non-zero. For simplicity, we use a normal prior density for *μ*, which is a conjugate prior for the normal density and so the posterior is also normal. Suppose that 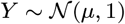 and 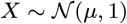. Let the prior for *Y* be 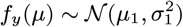 and the prior for *X* be 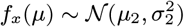. The “preferred prior” for use in generating the posterior based on both *X* and *Y* is 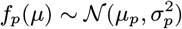. The posteriors are then

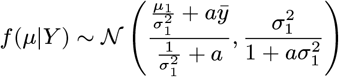

and

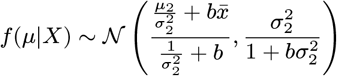

The approximation of the posterior of *μ*, given *X* and *Y*, is then

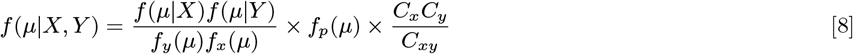

The true posterior is know in this case when *ρ* = 0. Let *Z* = *X* ∪ *Y* and *n* = *a* + *b*, then

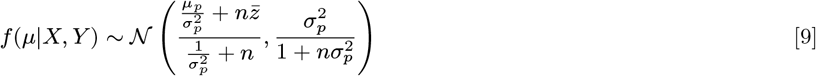

This simple case can be used to test methods for inferring the posterior from combined samples. Rather than doing MCMC, instead simply sample iid random variables from *f*(*μ*|*Y*), *f*(*μ*|*X*), *f_y_*(*μ*), and *f_x_*(*μ*) and use kernel density estimation to infer the density functions for each. Then apply equation 8 to estimate the posterior. The accuracy of the estimate can be determined by comparison with results from equation 9. For example, curves could be plotted for the true density versus the approximation. The approximate density will need to be renormalized so that it integrates to 1. The constant, *C*, to multiply values by to normalize could be estimated as

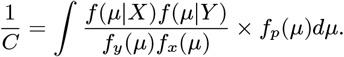

##### B. Numerical Issues Combining Posteriors

In a small number of cases, numerical issues arose when combining posteriors as using the packages described in the main text. In one case, no error messages resulted but the proportionality constant was on the order of 10^−12^. Likely due to numerical issues, this caused the mean latent integration time to be estimated to a value on the order of 10^6^, which has zero prior probability. This latent integration inference was removed from the analysis. Out of inferences for 90,000 latent integration times, 63 other analyses combining latent integration times from all four gene produced error messages related to non-integrable functions, and did not produced an estimate. This occurred for 8 latent times in the analysis of two genes only. These cases were removed from further analysis. These likely result when the posterior distributions from different genes are non-overlapping.

#### 9. Empirical Analysis of the Jones et al. dataset

Sequences originally published by Jones et al. (2018) were taken from GenBank (accession nos. MG822917-MG823179), and separated into patient 1 and patient 2 (13). The sequences from patient 1 were aligned using mafft (version 7.453) using the default settings (14). The sequences from patient 2 did not need to be aligned. The relative sample dates were determined using the collection date.

HIVtree was run with a burnin of 5,000 iterations, with 70,000 samples, sampling every other iteration. Two replicate MCMCs were run for each dataset. Convergence was checked by confirming no more than 5% of the mean internal nodes ages differed by more than 10 days between replicate MCMCs, 2.5% differed by more than 20 days, or any of the internal nodes differed by more than 100 days. Both pairs of MCMCs met this convergence criteria.

#### 10. Empirical Analysis of the Abrahams et al. dataset

Alignments for patients 217 and 257 originally published by Abrahams et al. (2019) were available from https://github.com/veg/ogv-dating/tree/master/results/alignments (15). There were multiple alignments for each data set and the “fasta_combined.msa” alignments were used. The week of sampling is included in the sequence name. Using the supplemental data table, the relative dates of sampling in units of days were determined. For some patients, there were multiple visit dates in the same week. In this case, the first visit date was used as the sample date for all sequences collected during that week. For each alignment, sequences were subsampled to include 10, 15, or 20 sequences from each pre-ART each collection time point and all outgrowth virus sequences. If less than the desired number of sequences were available at a given time point, all of the available sequences were used. While the sequences were aligned, some of the alignments had many gaps. Sites in the alignments were removed if they had more than 75%, 85%, or 95% gaps. Thus, for each of 8 starting alignments, 9 alignments were created. However, some of the alignments with gap removal were identical. Thus, a total of 46 unique alignments were created. HIVtree requires the sampling date to be at the end of the sequence name. Thus, the sequence names from the original publications were modified for our analyses.

Two replicate runs of HIVtree were run for each analysis. A burnin of 8,000 was used with samples taken every other iterations for a total of 80,000 samples. Thus, the MCMC was run for 168,000 iterations. Convergence of the MCMCs was checked by comparing the mean ages of the internal node ages. If more than 5% of the mean internal nodes ages differed by more than 10 days between replicate MCMCs, 2.5% differed by more than 20 days, or any of the internal nodes differed by more than 100 days, the MCMC was considered to not have converged. Two pairs of MCMCs did not converge. These were rerun with a a total of 150,000 samples, sampling every other iteration with a burnin of 8,000 iterations. Convergence was checked again with the same criteria as previously. Both pairs MCMCs had converged.

Each figure (S9 - S24) show the inferred integration date for each method, LR, LS, ML, and HIVtree. Each figure is for a single patient and gene. Some figures have two levels of gap removal instead of three because gap removal at different levels resulted in identical alignments. Thus, only the non-redundant results are shown. The gene names (e.g. ENV_4, NEF_1) match those in the original alignment names.

#### 11. Effect of the number of non-latent samples on method performance

The effect of tree size on the inference of latent samples was examined by changing the number of non-latent samples at each sample time. Using the simulated trees and alignments used in the main simulation analysis, the subsampling was changed from having 10 to 10, 15 or 20 non-latent sequence sampled every year for ten years. This results in a larger phylogenetic tree with the same number of latent sequences for each tree. Each tree was subsampled only one time for each number of non-latent sequences, rather than three times in the main analysis. The number of non-latent sequences at each sampling time does not have a large impact on bias (S25), MSE (S26), size of the 95% confidence intervals (S27), or the probability the inferred integration times fall within the 95% confidence intervals or credible sets (S28) for any of the methods.

As preliminary analysis did not show any trend with the other methods, this analysis was only run for the *p17* datasets with HIVtree. For the analyses with HIVtree, the priors were the same as in the main simulation analyses with HIVtree. The MCMCs were run with a burnin of 5,000 iterations, sampling every other iteration and sampling a total of 50,000 times. Two replicate MCMCs were run for each analysis. The difference between the mean times of the internal nodes was compared. The MCMCs were considered to have converged if this difference was no more than 10 days for at most 10% of the internal nodes, 20 days for at most 5% of the internal nodes, and no more than 100 days for any of the internal nodes. 10 pairs of MCMCs did not converge. These were run again with a burnin of 10,000 iterations, sampling 100,000 times with sampling every other iteration. The above convergence criteria were checked again. All MCMCs were considered to have converged.

**Fig. S1.**
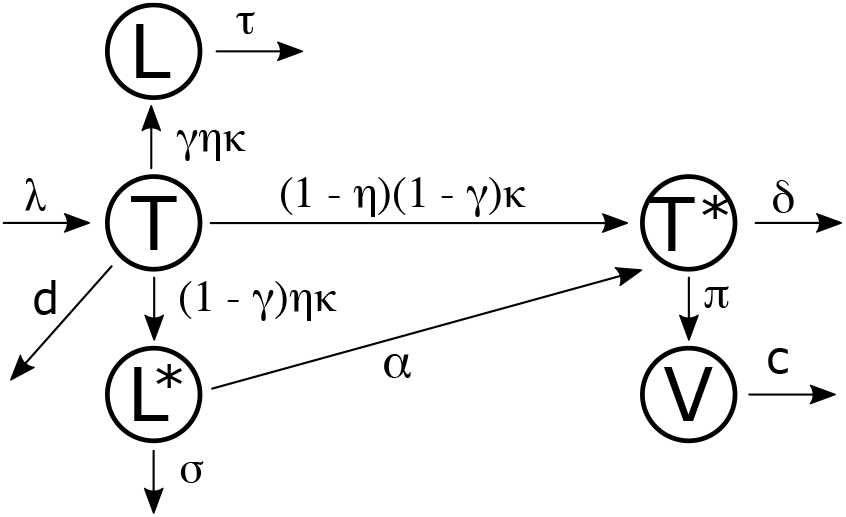
Within-host viral dynamics model

**Fig. S2.**
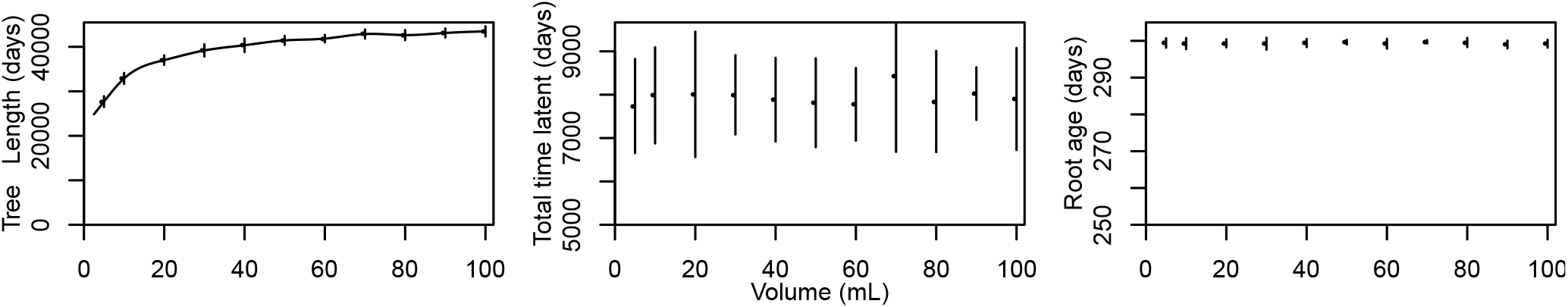
Impact of simulation volume on properties of genealogies. 50 active and 20 latent viruses were sampled at 75, 100, 200, and 300 days. 10 simulations were run for each simulation volume. Other simulation parameters match those in S1. Standard error is shown.

**Table S1.** Simulation parameters. The parameters from (2) are for patient 7. *κ* is typically estimated as the rate constant of new infections of replication-competent cells, which is *κ*(1 – *γ*)(1 – *η*) in this model. Thus, the empirical estimates of *κ*, as presented in the table, is divided by (1 – *γ*)(1 – *η*) to obtain the parameter value used in the model.

| Parameter | Description | Value |
| --- | --- | --- |
| $\lambda$ | Birth rate of uninfected cells | $170 \frac{\text{cell}}{\text{mL} \times \text{day}}$ (2) |
| $d$ | Death rate of uninfected cells | $0.017 \frac{1}{\text{day}}$ (2) |
| $\kappa$ | Transition rate from uninfected to actively infected cells | $8.0 \times 10^{-7} \frac{\text{mL}}{\text{virion} \times \text{day}}$ (2) |
| $\delta$ | Death rate of actively infected cells | $0.31 \frac{1}{\text{day}}$ (2) |
| $\pi$ | Viral birth rate | $730 \frac{\text{virions}}{\text{cell day}}$ (2) |
| $c$ | Viral clearance rate | $3 \frac{1}{\text{day}}$ (2) |
| $\eta$ | Proportion of newly infected cells that are latent | $1.16 \times 10^{-3}$ (3) |
| $\alpha$ | Rate of activation of replication-competent, latent cells | $5.7 \times 10^{-5} \frac{1}{\text{day}}$ (16, 17) |
| $\gamma$ | Proportion of viruses that are defective | 0.95 (18) |
| $\sigma$ | Death rate of latent, replication-competent cells | $5.2 \times 10^{-4} \frac{1}{\text{day}}$ (19) |
| $\tau$ | Death rate of latent, replication-incompetent cells | $1.1 \times 10^{-4} \frac{1}{\text{day}}$ (19) |

**Fig. S3.**
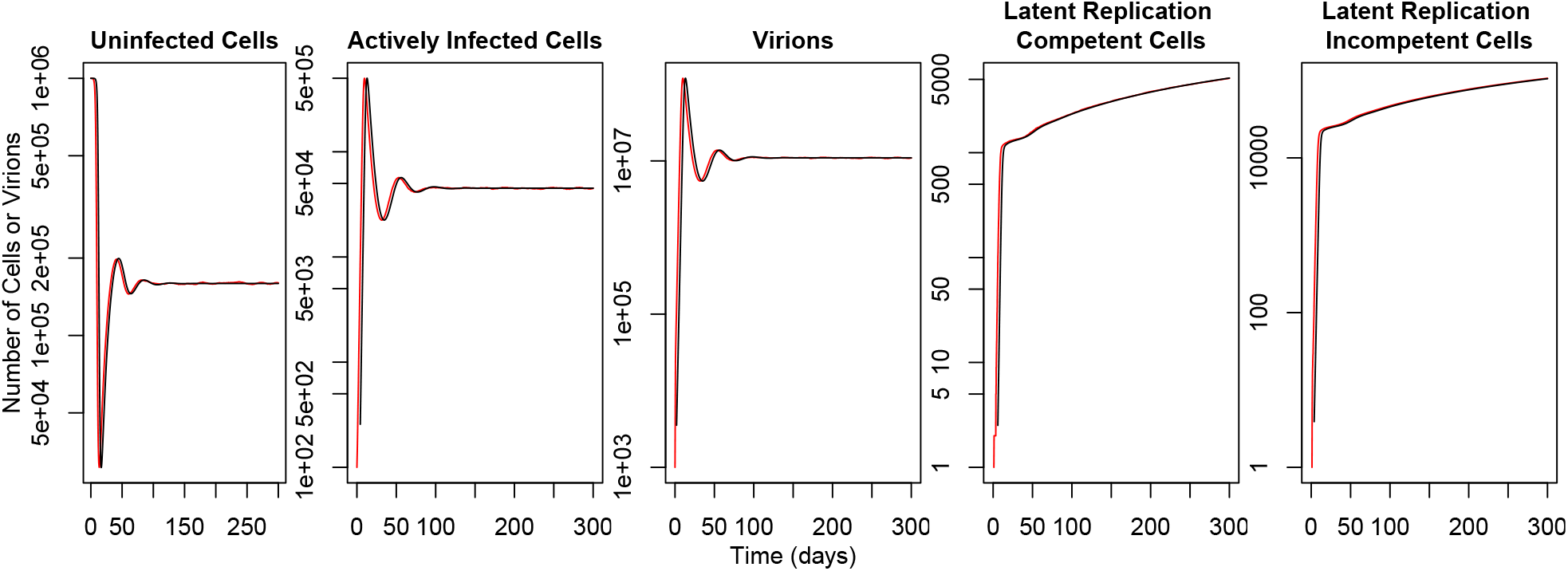
Predicted population sizes in the deterministic model and observed population sizes in the stochastic model are very similar. For both models, a blood volume of 10mL was modeled using the parameters listed in Table S1. The initial population sizes are 10^4^ target cells/mL, 1 actively infected cell/mL, and 10 virions/mL. The deterministic model is shown in black, and one realization of the stochastic simulation is shown in red. In comparison to the initial conditions described in the text, a larger number of actively infected cells was used to limit the stochastic effects of small population sizes, allowing for a comparison when the virus is unlikely to become extinct.

**Table S2.** DNA simulation parameters. *μ* is in units of expected number of substitutions per day per base. The genes simulated do not cover the entire genes.

| Region | HXB2 start | HXB2 end | $\mu$ | $\alpha$ | $\kappa$ | $\pi_A$ | $\pi_C$ | $\pi_G$ | $\pi_T$ |
| --- | --- | --- | --- | --- | --- | --- | --- | --- | --- |
| C1V2 | 6213 | 7037 | $3.56 \times 10^{-5}$ | 0.4294 | 6.9801 | 0.35322 | 0.17636 | 0.21123 | 0.259191 |
| nef | 8797 | 9414 | $1.34 \times 10^{-5}$ | 0.4878 | 8.9138 | 0.30641 | 0.21240 | 0.28265 | 0.19853 |
| p17 | 817 | 1207 | $8.9 \times 10^{-6}$ | 0.5306 | 10.6361 | 0.39393 | 0.18392 | 0.25040 | 0.17175 |
| tat | 5831 | 5962 | $9.9 \times 10^{-6}$ | 0.7283 | 7.1751 | 0.29841 | 0.21021 | 0.23449 | 0.25689 |

**Fig. S4.**
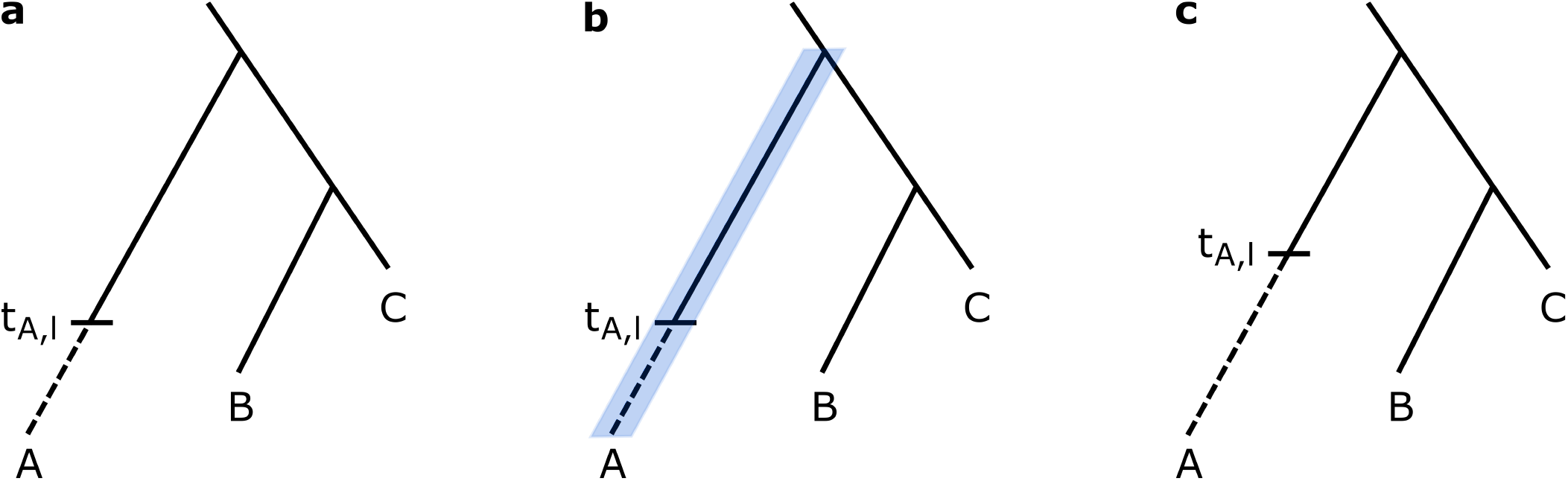
Proposal steps in the MCMC for latency times. Tips B and C correspond to non-latent sequences. At some time in the past, *t_A,l_*, lineage A became latent. The dashed line shows when the lineages was latent. (a) Starting from the current latent time, (b) a new time can be proposed anywhere between the sample time and the age of the parent node, shown in blue. (c) Once a time is proposed, the move can be accepted or rejected. In this case, the move is accepted and the time is updated. For the calculation of the likelihood, the branch lengths correspond to the length of the solid lines only.

**Fig. S5.**
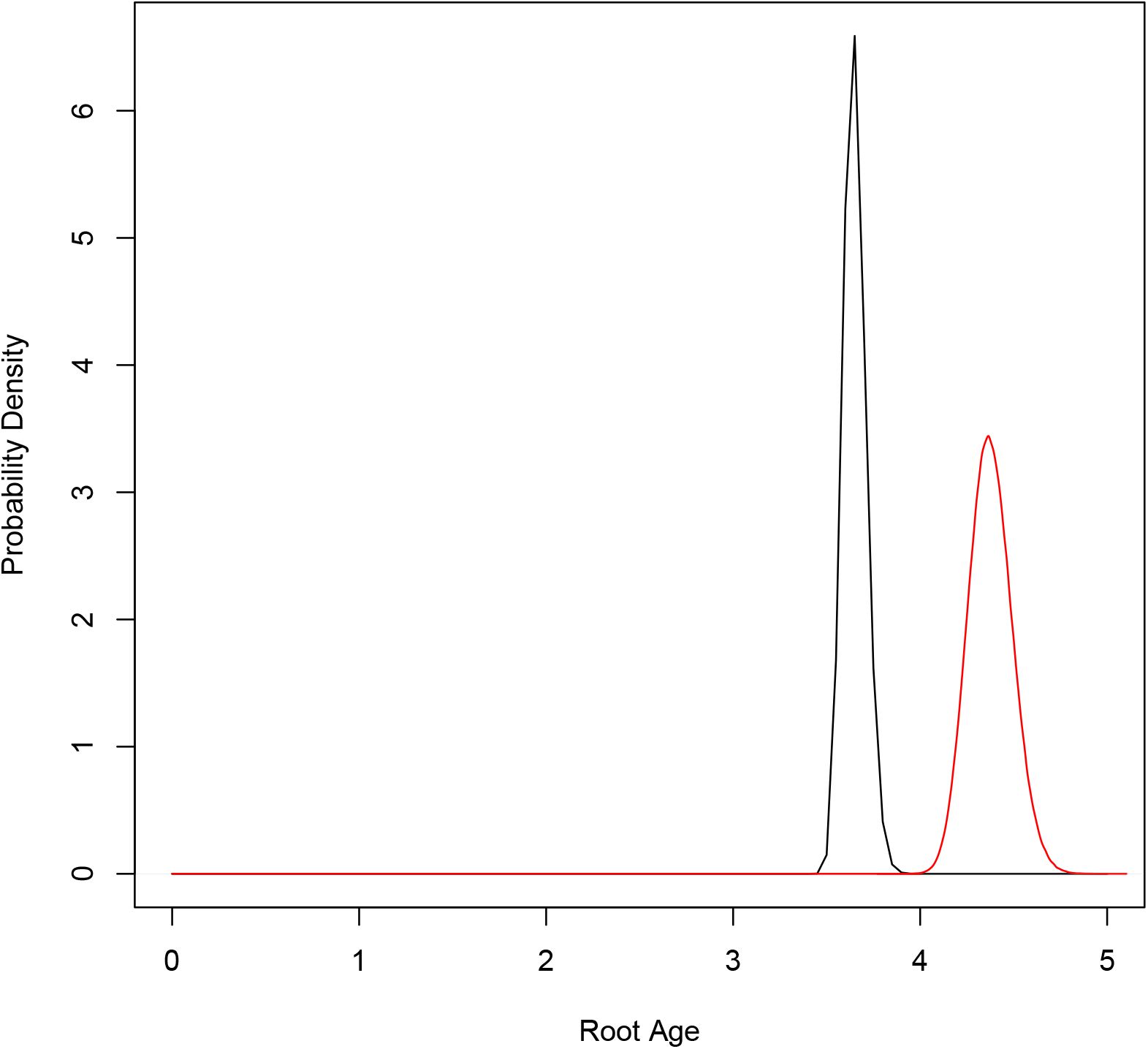
The user input prior for the root age is not the same as the prior determined by running HIVtree without data. The black line shows the user input root age prior of Gamma(36.5, 100) on a tree with a last sample time of 3285 days before present with a time unit of 1000. This gives as mean root age of 3.65 in the time units used in HIVtree. This is the same as all of the simulated trees in our analyses. Using a simulated dataset for C1V2, a tree toplogy was inferred with RAxML and outgroup rooted. This tree was used to run HIVtree under the prior. The red line shows the results, in which the root age is older than the user input prior.

**Fig. S6.**
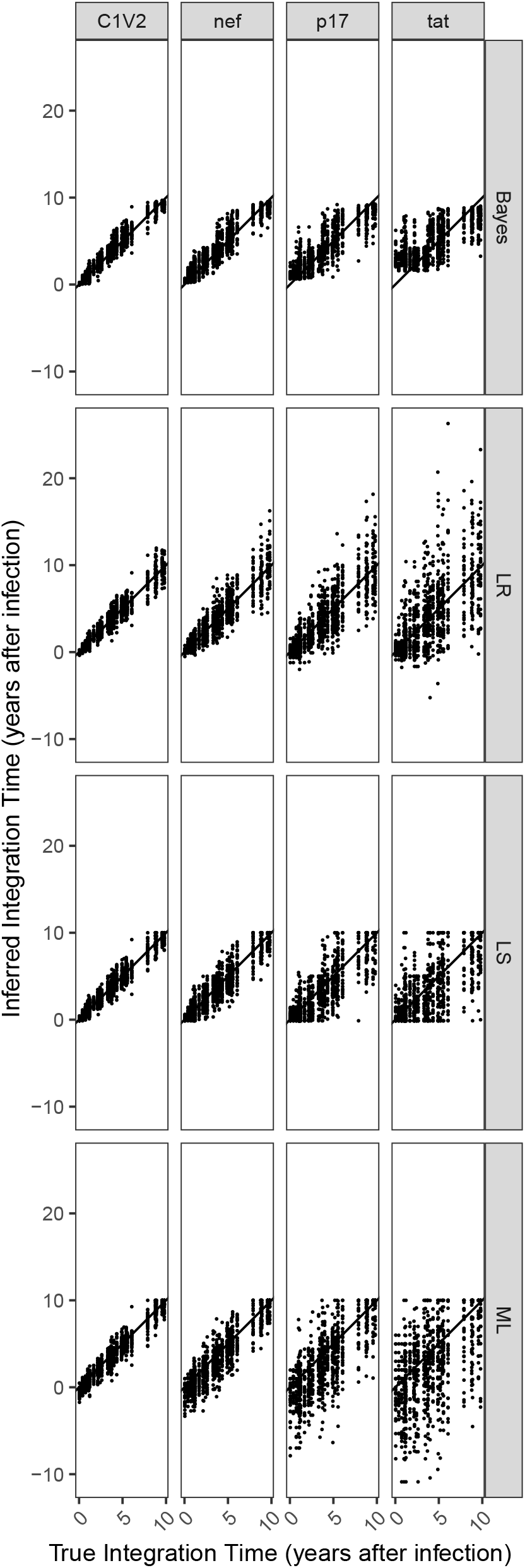
For a fixed tree toplogy, there are 30 latent integration times for each of the 30 alignments for a given gene. The line has slope 1 and intercept 0.

**Fig. S7.**
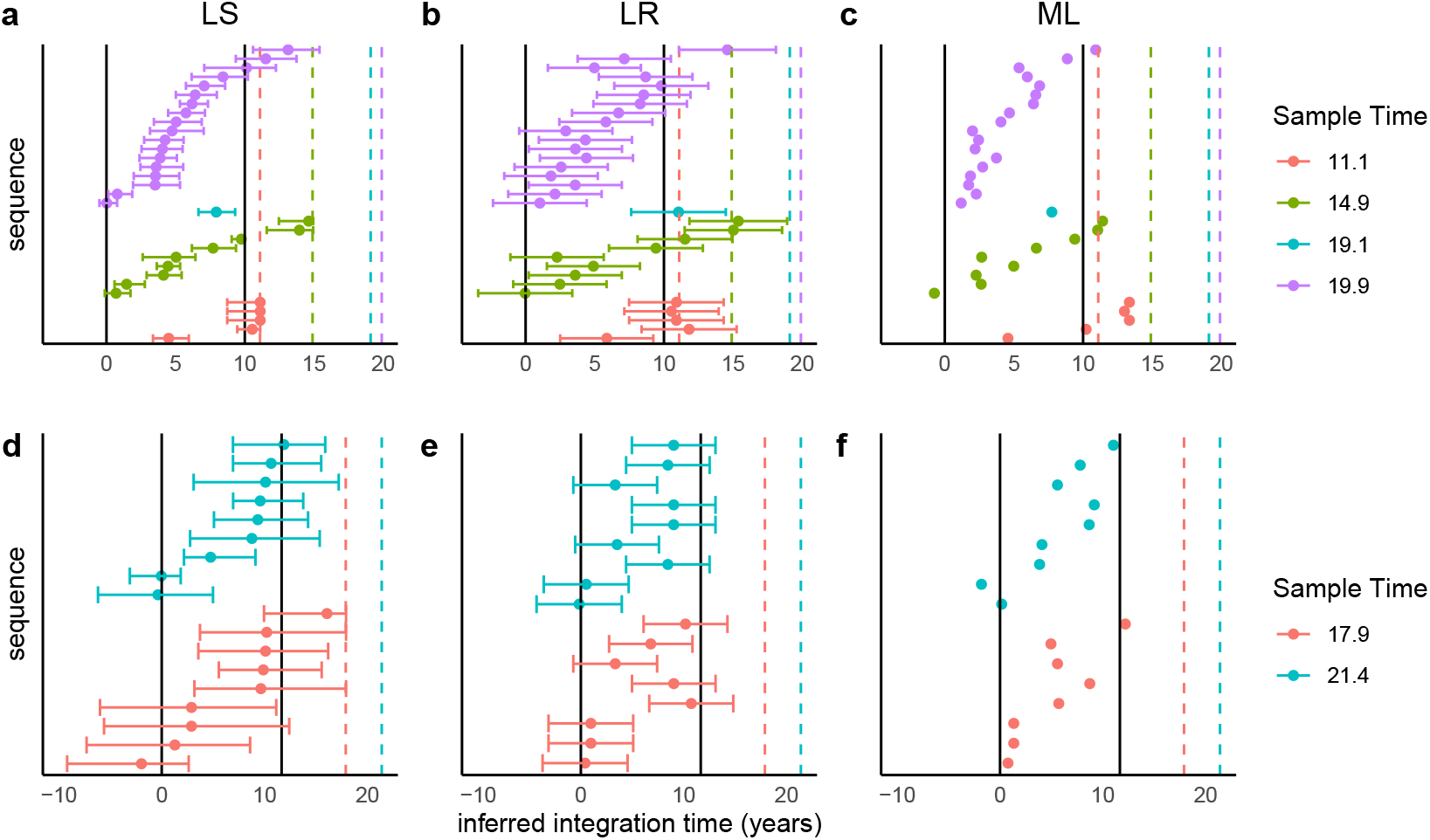
(a-c) and (d-f) show the inferred integration dates for each sequence from patient 1 and 2, respectively. (a,d), (b,e), and (c,f) show inferences from LS, LR, and ML, respectively. The vertical lines show the first positive date (left) and start of cART (right). The bar show 95% confidence intervals for LS and LR. Confidence intervals are not inferred in the ML method. With sample time 11.1 for patient 1, three of the latent integration times inferred with ML and one with LR are after the sampling date. The LS method is bounded at the sample time, but those sequences are inferred to have been integrated at the sample time.

**Fig. S8.**
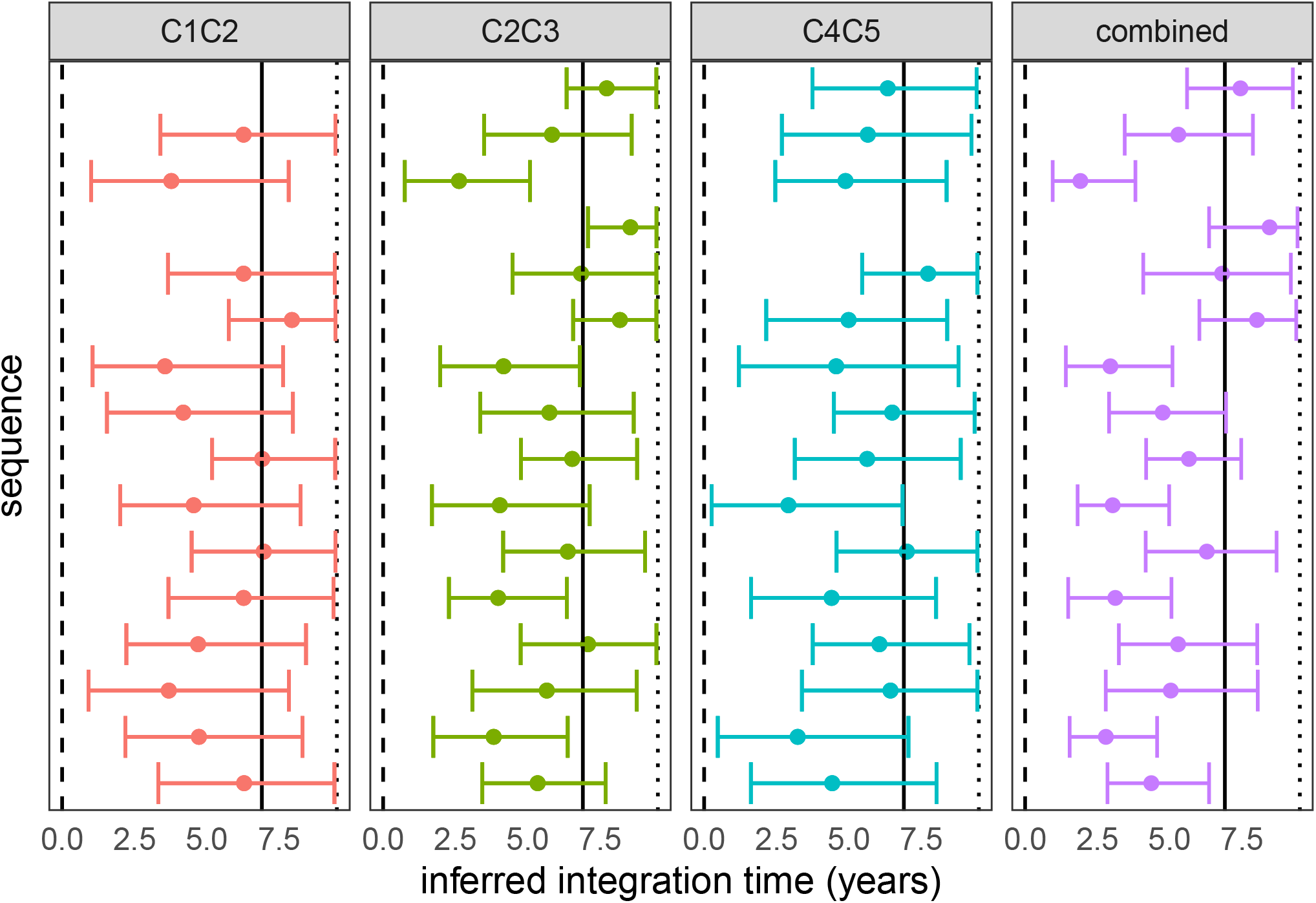
Each of the left three panels shows the integration times inferred using HIVtree for a single sequence. The panel on the right shows the inferred integration times when the posterior estimate for the three sequences are combined. The results are from patient 217 (15). 10 non-latent sequence were used as each available timepoint and sites with more than 75% gaps were removed from the alignment prior to analysis, as described in SI section 10. The dashed line shows the infection time, the solid line shows the start of ART, and the dotted line shows the sample time.

**Fig. S9.**
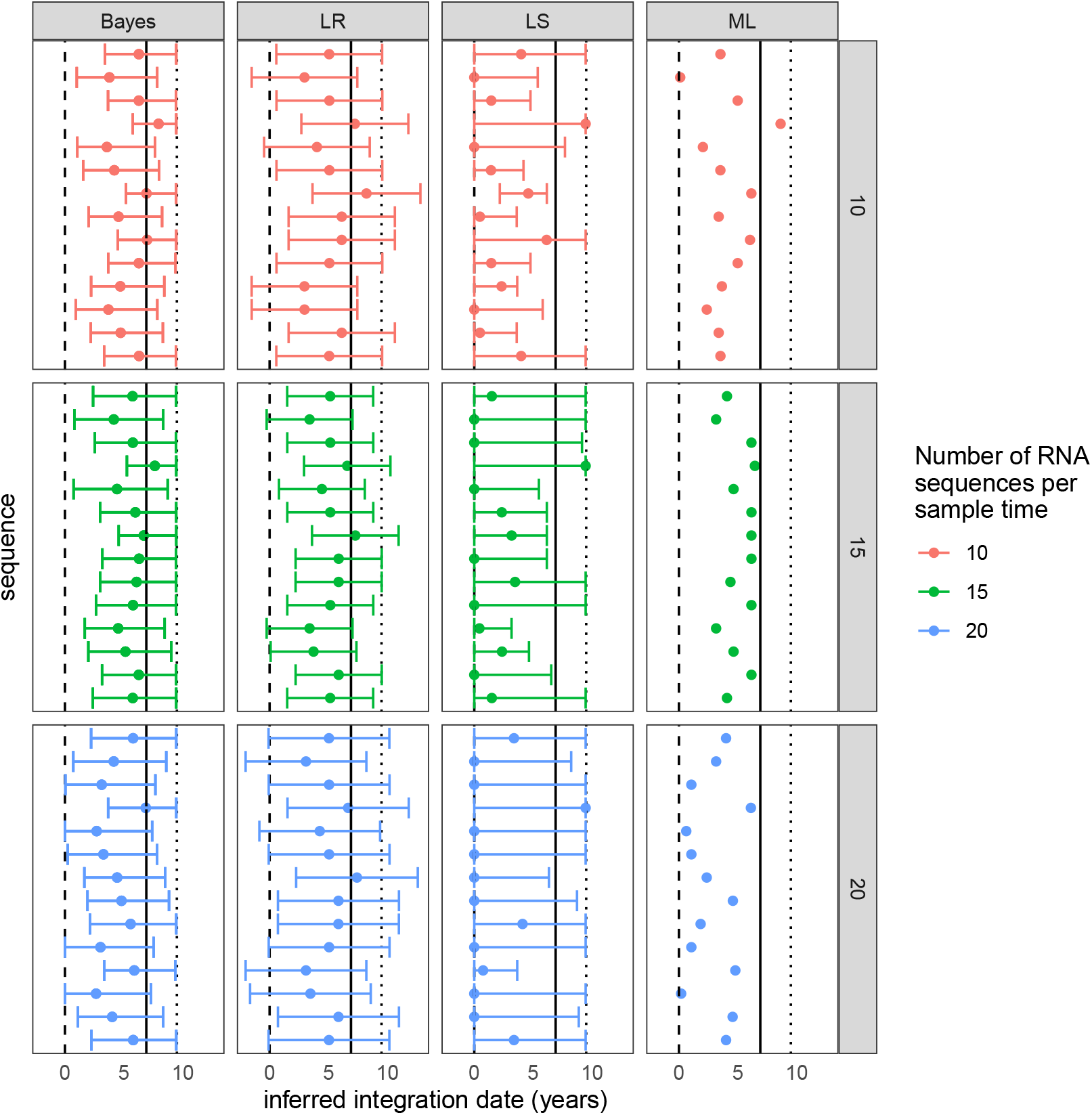
The inferred latent integration dates for Env_2 from patient 217 are shown for each method. 95% confidence intervals are shown for the LR and LS methods, and the 95% credible interval is shown for HIVTree. Sequences are shown in the same order in each panel. The vertical lines show the time of infection (dashed), time of treatment start (solid) and the time of sampling (dotted). The color shows the number of RNA sequences subsampled from the original alignment at each sample time. If fewer sequences were available then the number indicated by the color at a given time, all available sequences were used. Sites with greater than 75% missing gaps have been removed from the alignment.

**Fig. S10.**
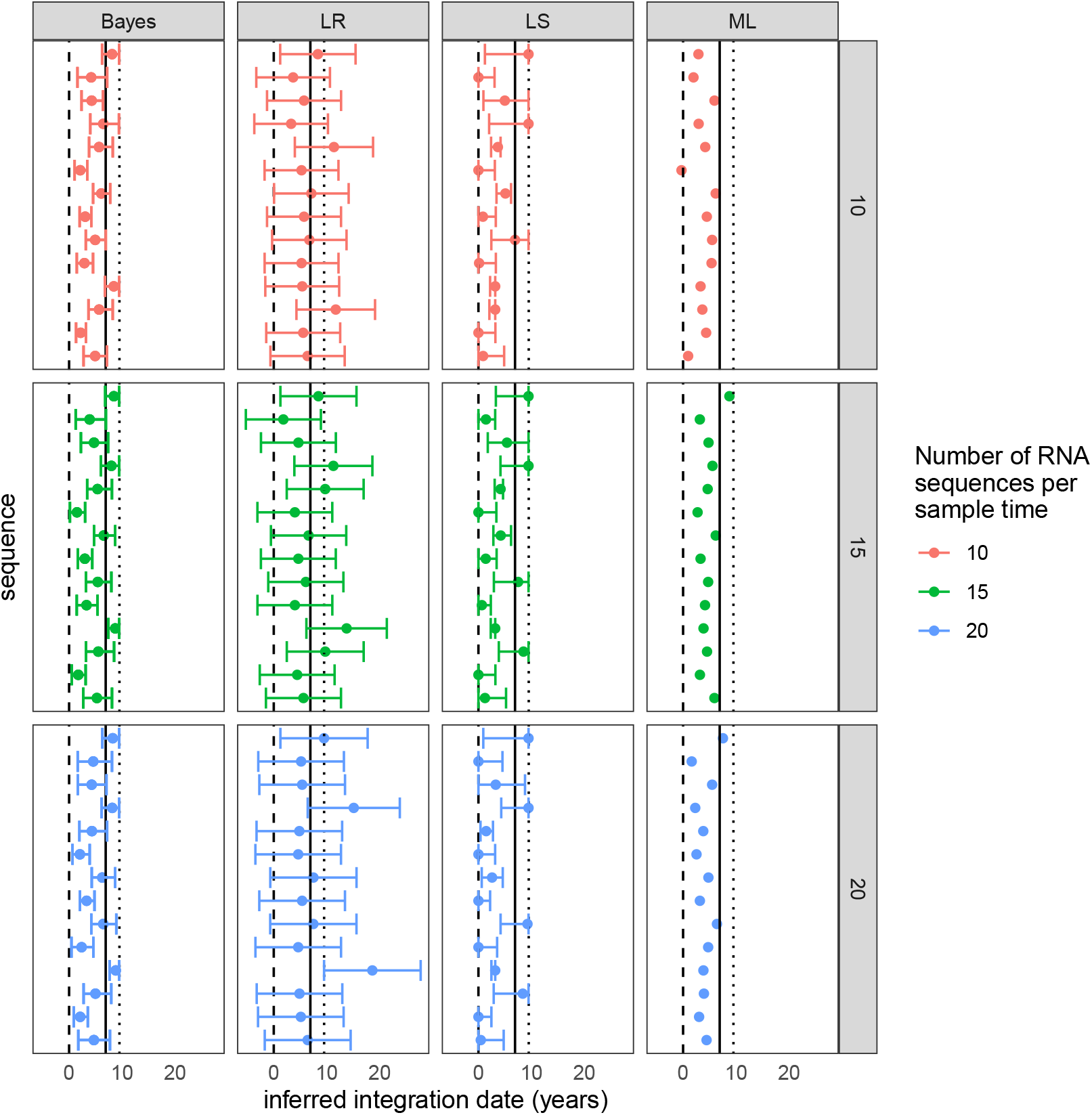
The inferred latent integration dates for Env_2 from patient 217 are shown for each method. 95% confidence intervals are shown for the LR and LS methods, and the 95% credible interval is shown for HIVTree. Sequences are shown in the same order in each panel. The vertical lines show the time of infection (dashed), time of treatment start (solid) and the time of sampling (dotted). The color shows the number of RNA sequences subsampled from the original alignment at each sample time. If fewer sequences were available then the number indicated by the color at a given time, all available sequences were used. Sites with greater than 95% missing gaps have been removed from the alignment.

**Fig. S11.**
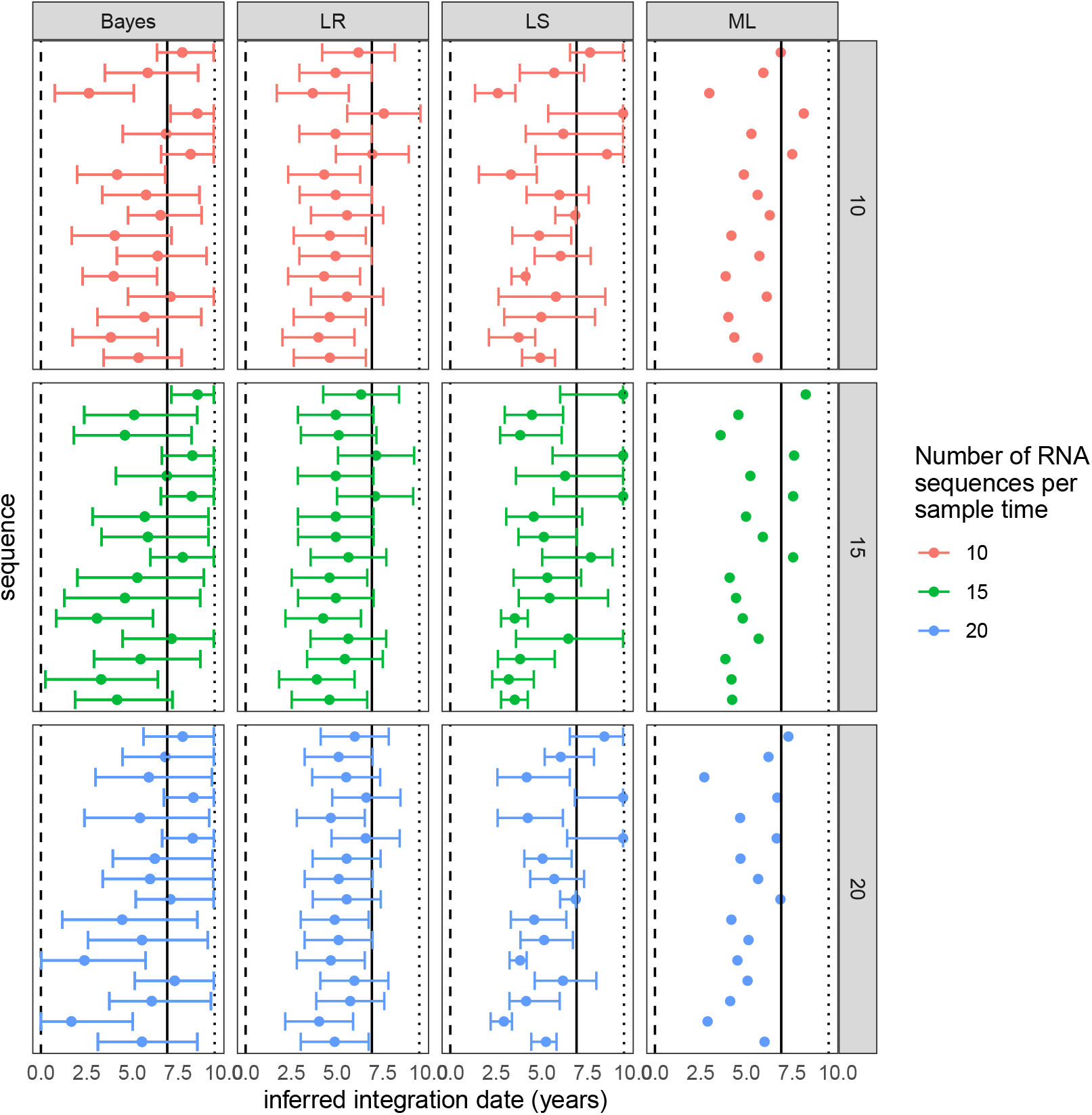
The inferred latent integration dates for Env_3 from patient 217 are shown for each method. 95% confidence intervals are shown for the LR and LS methods, and the 95% credible interval is shown for HIVTree. Sequences are shown in the same order in each panel. The vertical lines show the time of infection (dashed), time of treatment start (solid) and the time of sampling (dotted). The color shows the number of RNA sequences subsampled from the original alignment at each sample time. If fewer sequences were available then the number indicated by the color at a given time, all available sequences were used. Sites with greater than 75% missing gaps have been removed from the alignment.

**Fig. S12.**
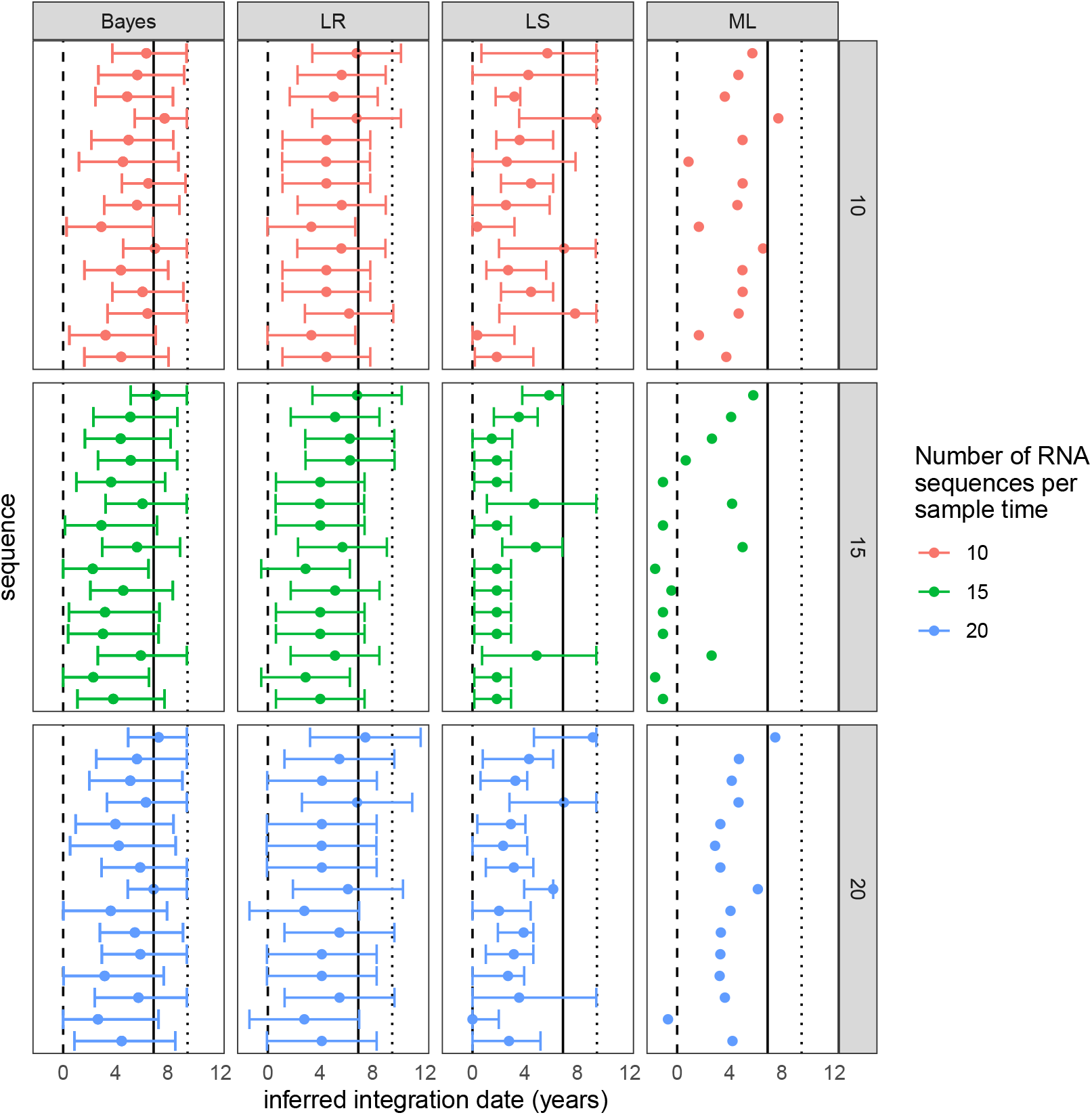
The inferred latent integration dates for Env_4 from patient 217 are shown for each method. 95% confidence intervals are shown for the LR and LS methods, and the 95% credible interval is shown for HIVTree. Sequences are shown in the same order in each panel. The vertical lines show the time of infection (dashed), time of treatment start (solid) and the time of sampling (dotted). The color shows the number of RNA sequences subsampled from the original alignment at each sample time. If fewer sequences were available then the number indicated by the color at a given time, all available sequences were used. Sites with greater than 75% missing gaps have been removed from the alignment.

**Fig. S13.**
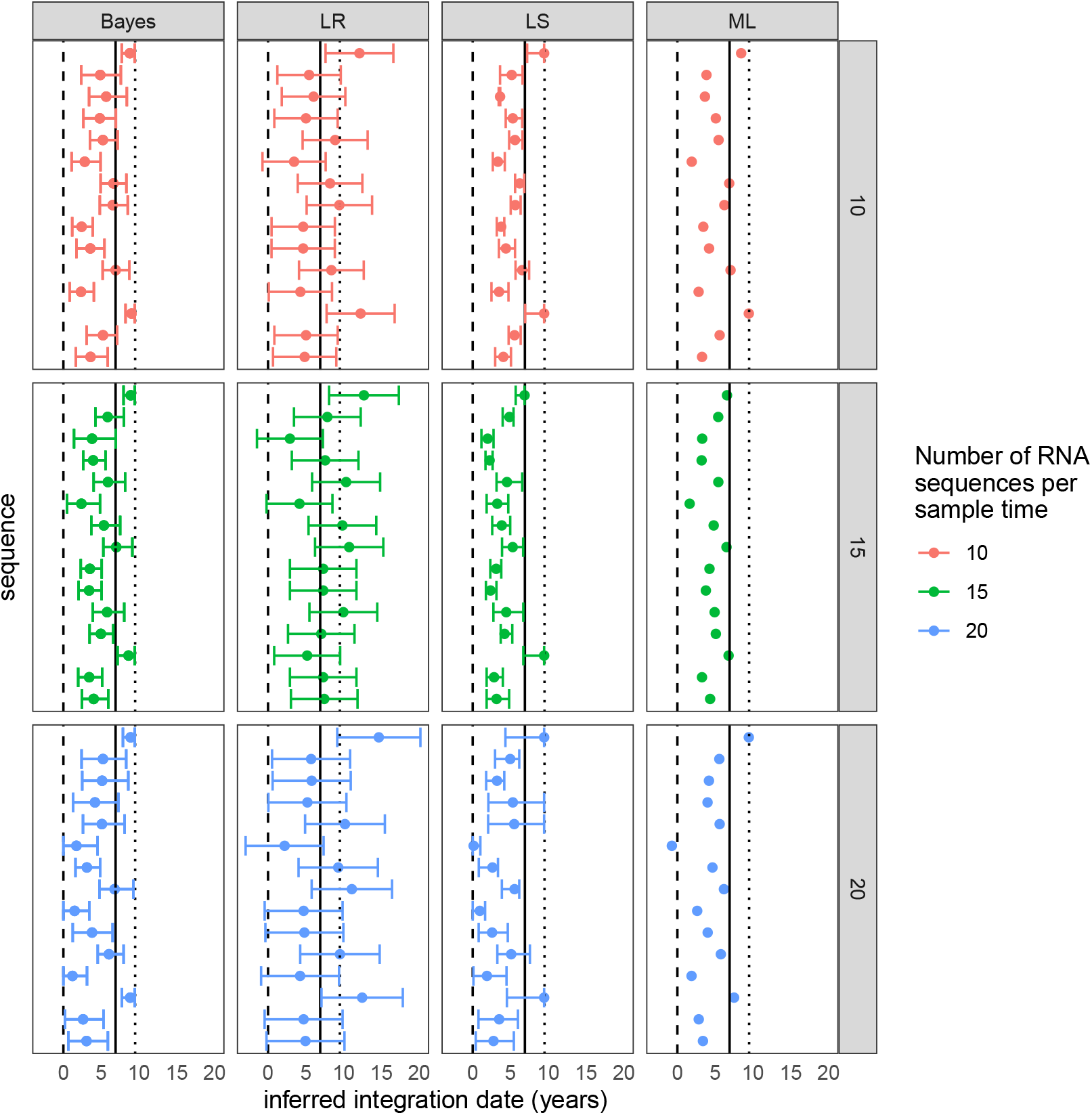
The inferred latent integration dates for Env_4 from patient 217 are shown for each method. 95% confidence intervals are shown for the LR and LS methods, and the 95% credible interval is shown for HIVTree. Sequences are shown in the same order in each panel. The vertical lines show the time of infection (dashed), time of treatment start (solid) and the time of sampling (dotted). The color shows the number of RNA sequences subsampled from the original alignment at each sample time. If fewer sequences were available then the number indicated by the color at a given time, all available sequences were used. Sites with greater than 95% missing gaps have been removed from the alignment.

**Fig. S14.**
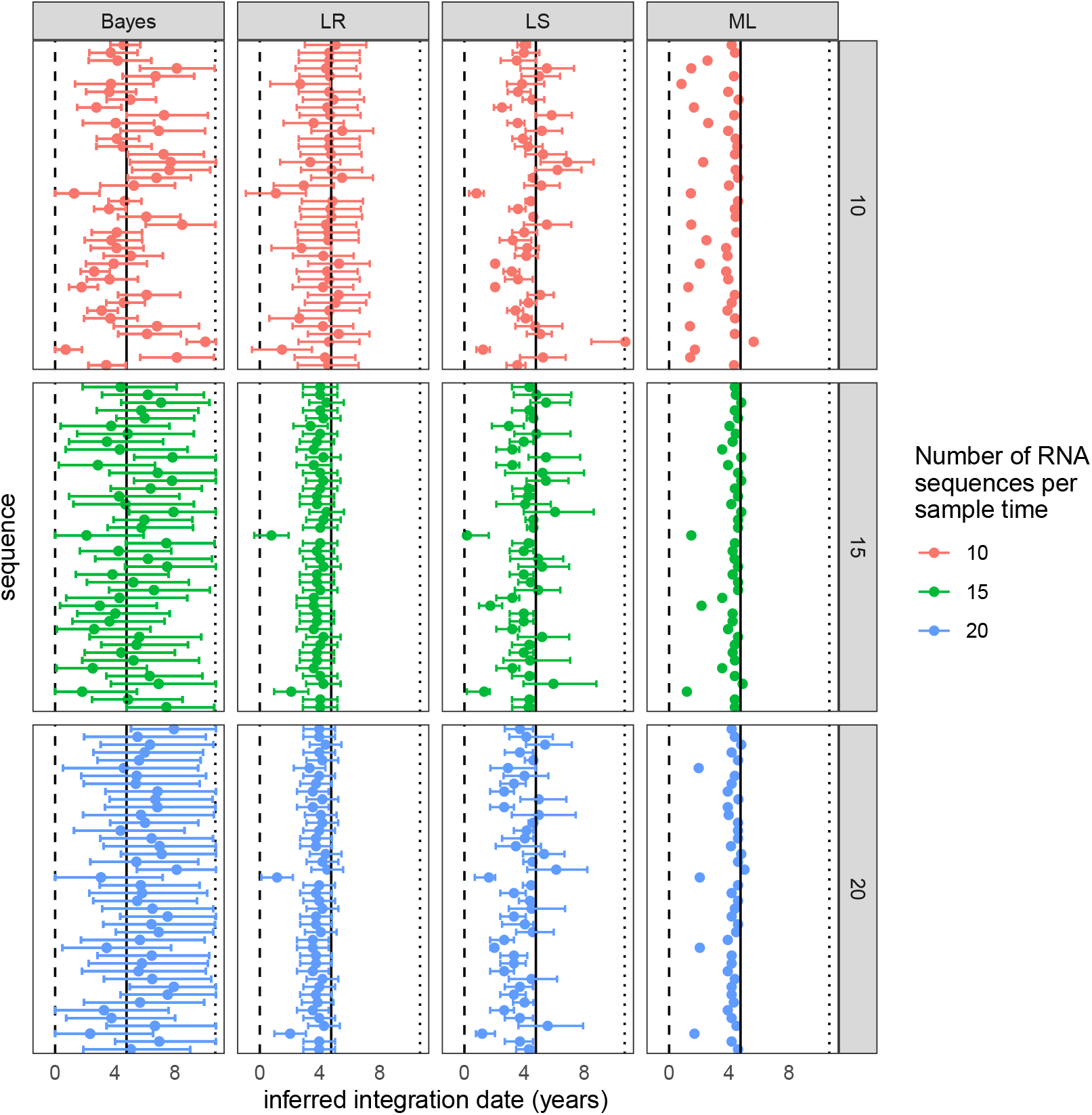
The inferred latent integration dates for Env_2 from patient 257 are shown for each method. 95% confidence intervals are shown for the LR and LS methods, and the 95% credible interval is shown for HIVTree. Sequences are shown in the same order in each panel. The vertical lines show the time of infection (dashed), time of treatment start (solid) and the time of sampling (dotted). The color shows the number of RNA sequences subsampled from the original alignment at each sample time. If fewer sequences were available then the number indicated by the color at a given time, all available sequences were used. Sites with greater than 75% missing gaps have been removed from the alignment.

**Fig. S15.**
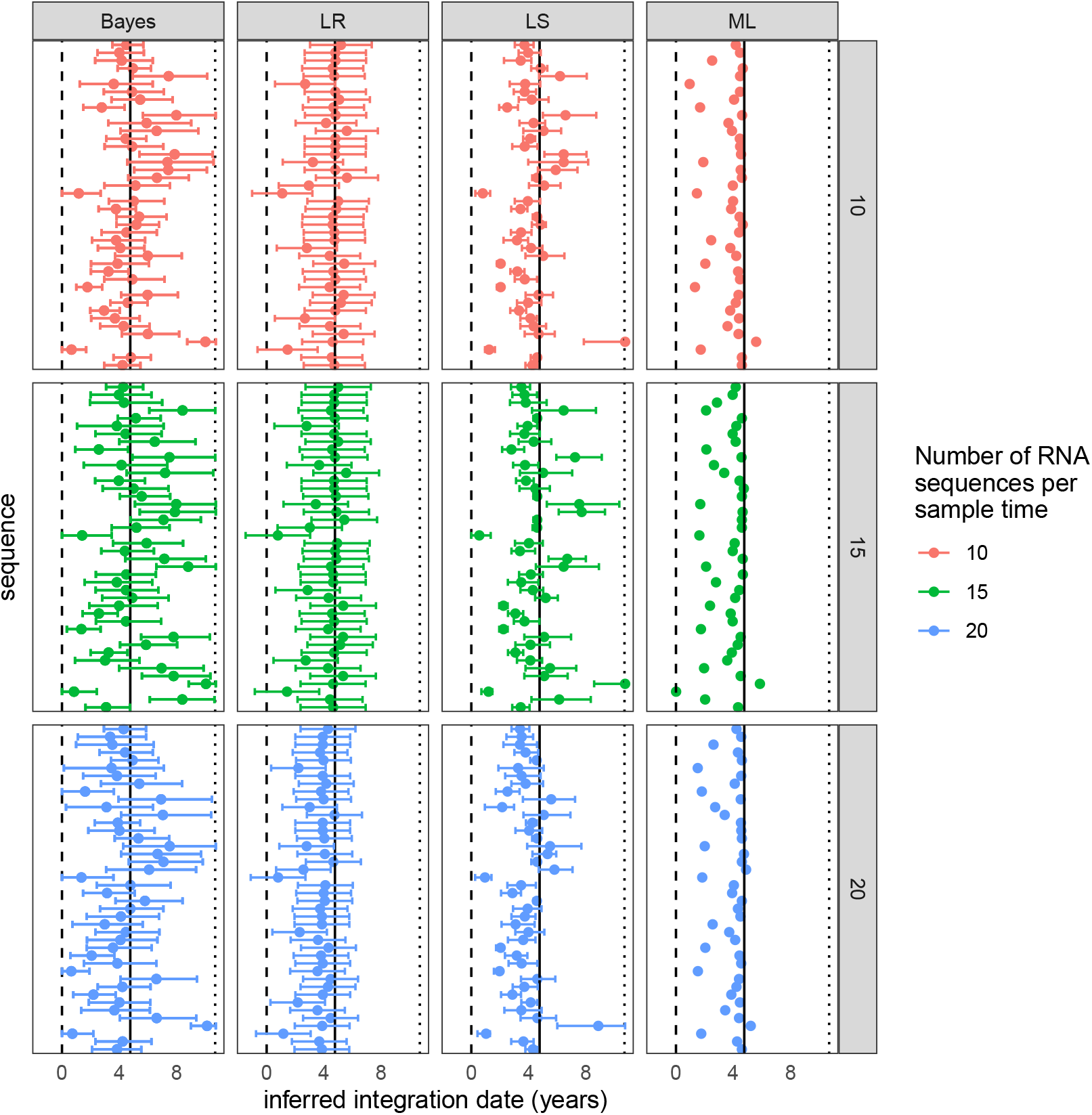
The inferred latent integration dates for Env_2 from patient 257 are shown for each method. 95% confidence intervals are shown for the LR and LS methods, and the 95% credible interval is shown for HIVTree. Sequences are shown in the same order in each panel. The vertical lines show the time of infection (dashed), time of treatment start (solid) and the time of sampling (dotted). The color shows the number of RNA sequences subsampled from the original alignment at each sample time. If fewer sequences were available then the number indicated by the color at a given time, all available sequences were used. Sites with greater than 85% missing gaps have been removed from the alignment.

**Fig. S16.**
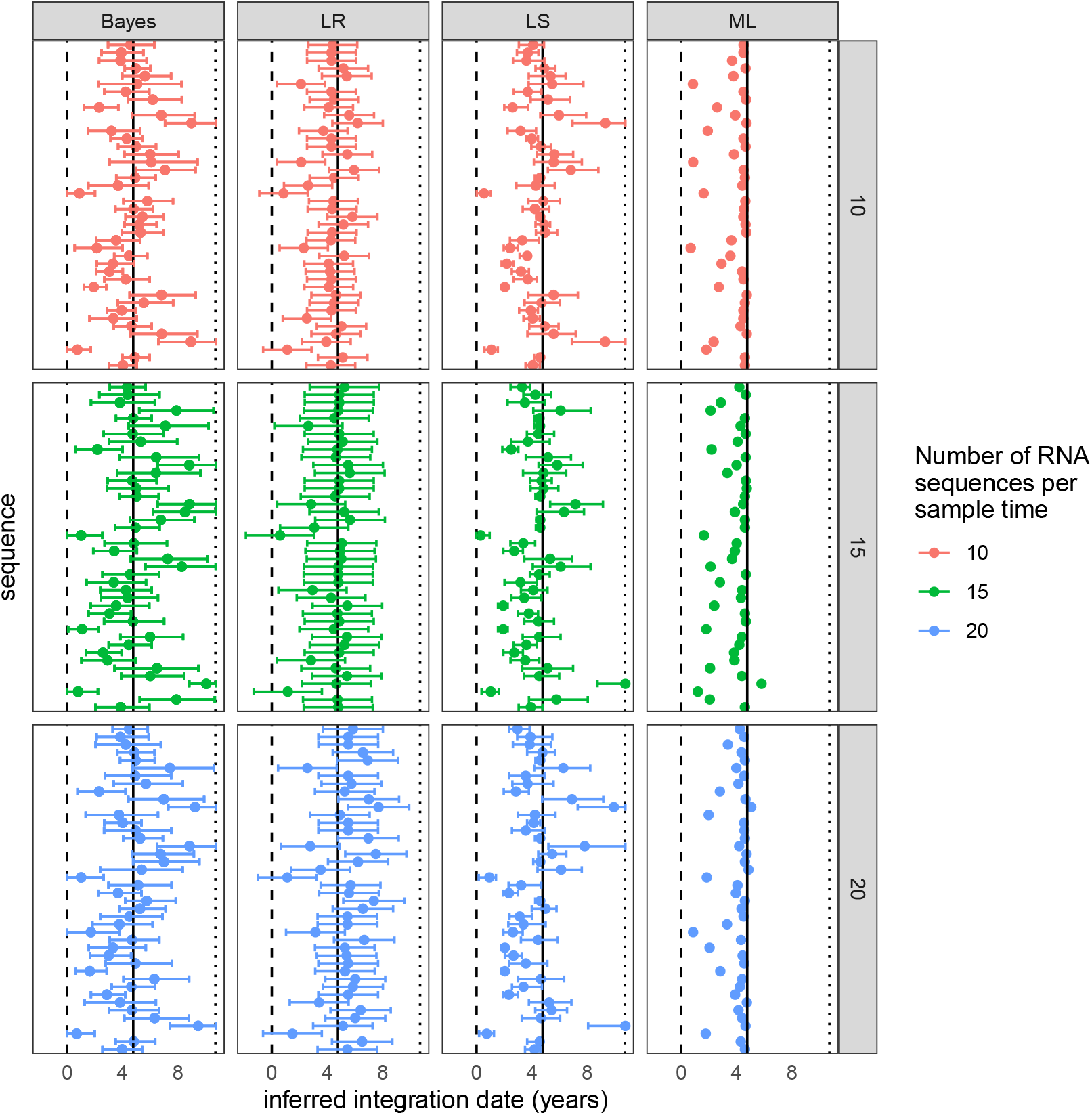
The inferred latent integration dates for Env_2 from patient 257 are shown for each method. 95% confidence intervals are shown for the LR and LS methods, and the 95% credible interval is shown for HIVTree. Sequences are shown in the same order in each panel. The vertical lines show the time of infection (dashed), time of treatment start (solid) and the time of sampling (dotted). The color shows the number of RNA sequences subsampled from the original alignment at each sample time. If fewer sequences were available then the number indicated by the color at a given time, all available sequences were used. Sites with greater than 95% missing gaps have been removed from the alignment.

**Fig. S17.**
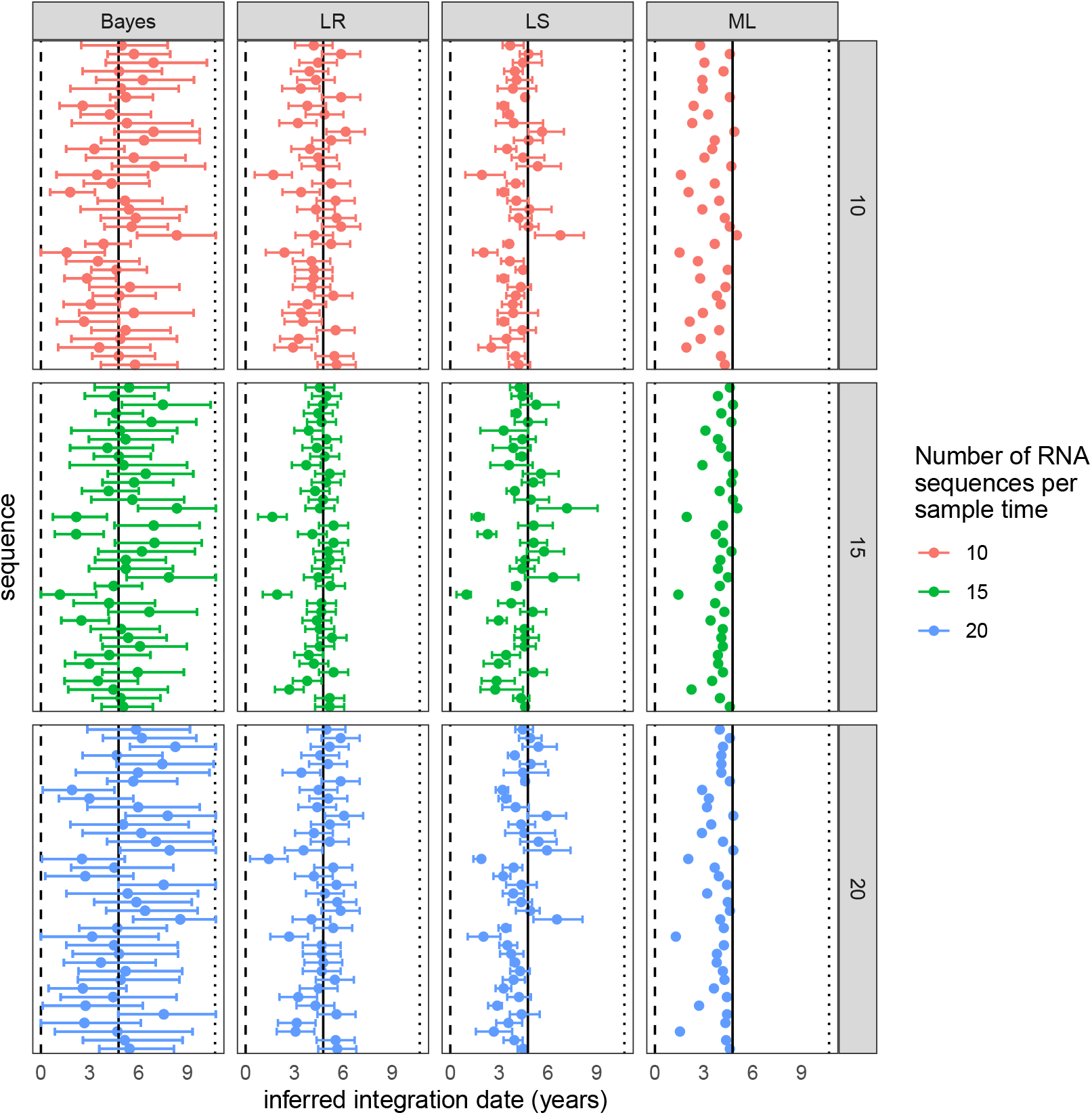
The inferred latent integration dates for Env_3 from patient 257 are shown for each method. 95% confidence intervals are shown for the LR and LS methods, and the 95% credible interval is shown for HIVTree. Sequences are shown in the same order in each panel. The vertical lines show the time of infection (dashed), time of treatment start (solid) and the time of sampling (dotted). The color shows the number of RNA sequences subsampled from the original alignment at each sample time. If fewer sequences were available then the number indicated by the color at a given time, all available sequences were used. Sites with greater than 75% missing gaps have been removed from the alignment.

**Fig. S18.**
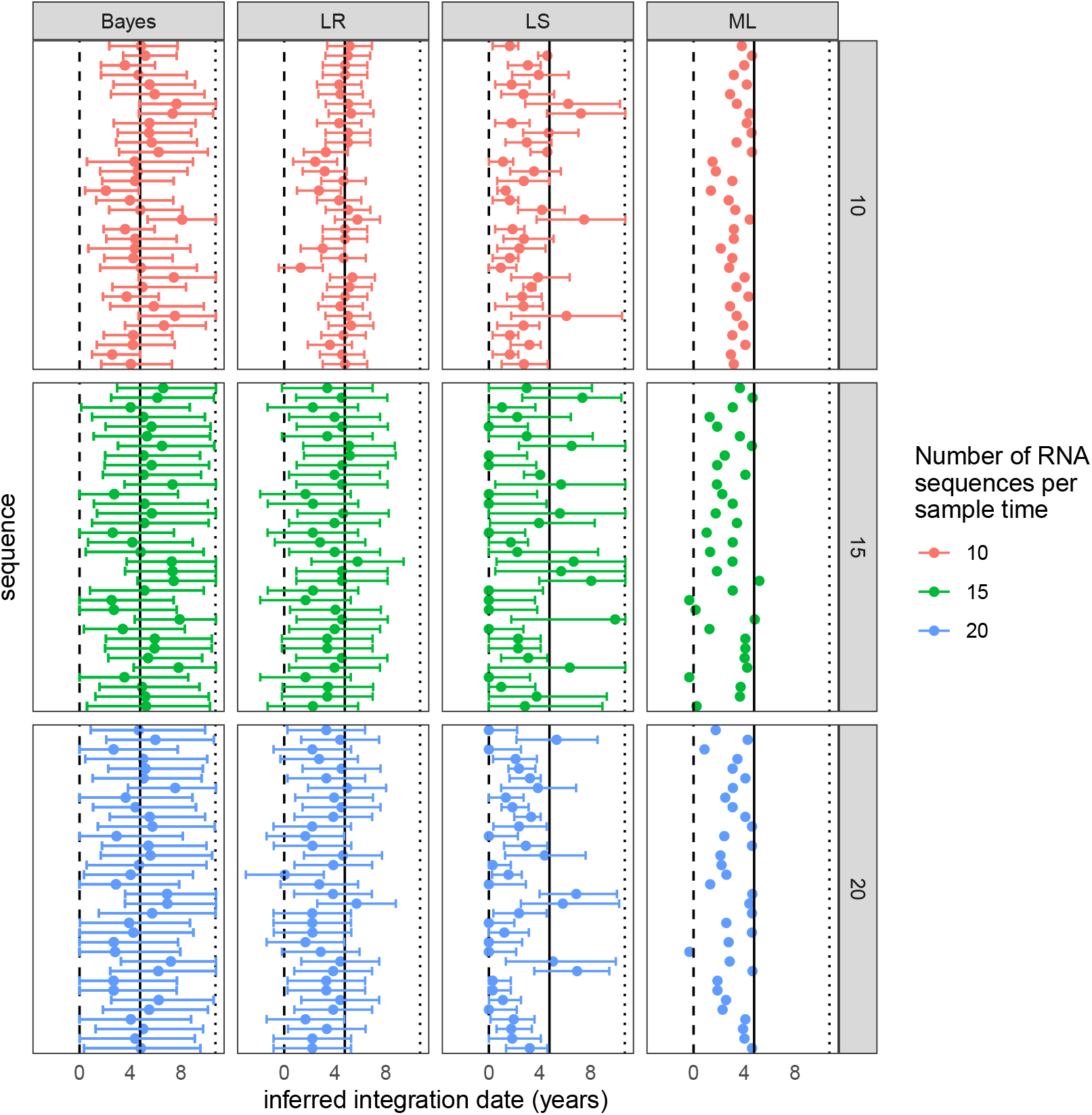
The inferred latent integration dates for Env_4 from patient 257 are shown for each method. 95% confidence intervals are shown for the LR and LS methods, and the 95% credible interval is shown for HIVTree. Sequences are shown in the same order in each panel. The vertical lines show the time of infection (dashed), time of treatment start (solid) and the time of sampling (dotted). The color shows the number of RNA sequences subsampled from the original alignment at each sample time. If fewer sequences were available then the number indicated by the color at a given time, all available sequences were used. Sites with greater than 75% missing gaps have been removed from the alignment.

**Fig. S19.**
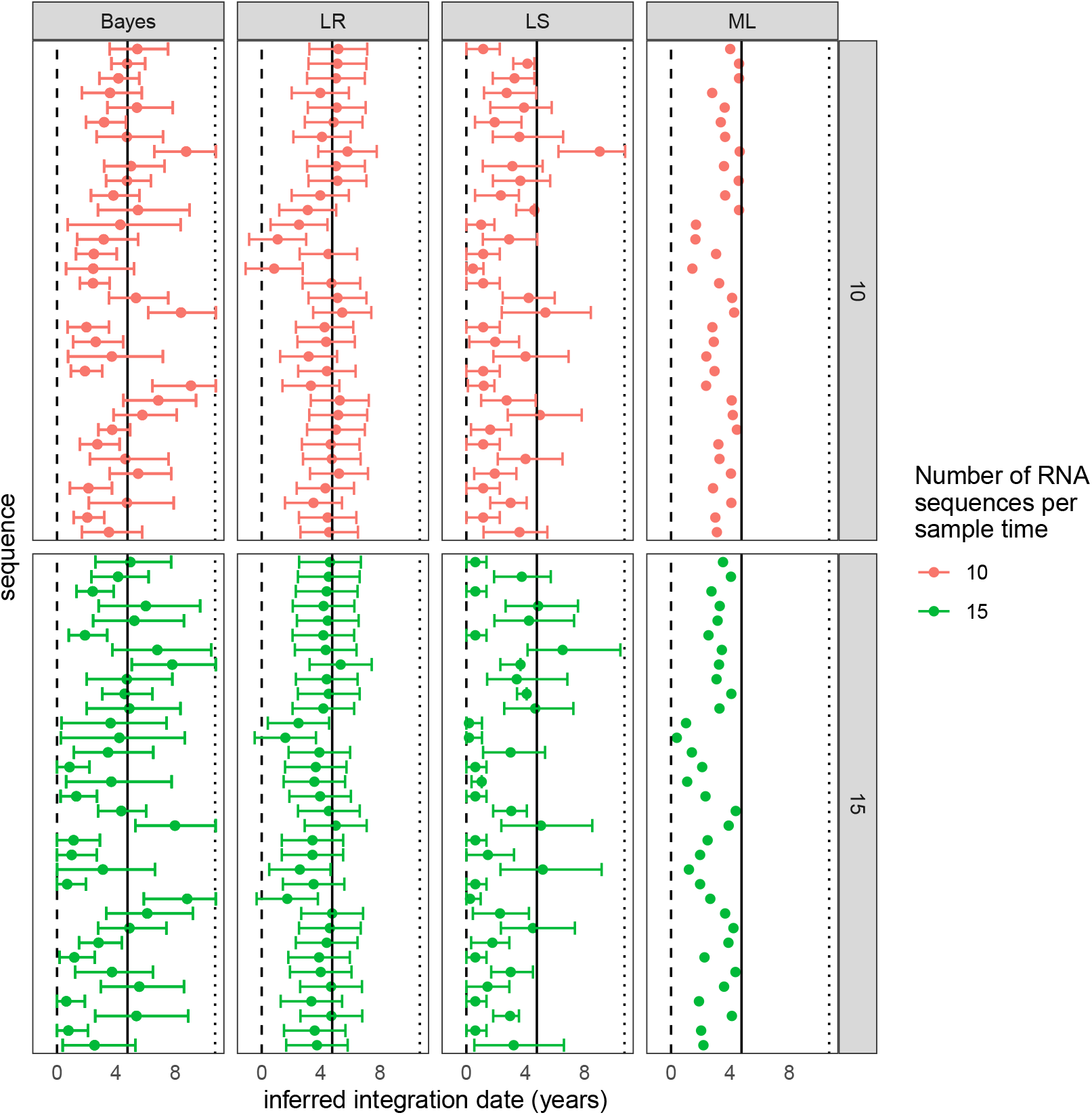
The inferred latent integration dates for Env_4 from patient 257 are shown for each method. 95% confidence intervals are shown for the LR and LS methods, and the 95% credible interval is shown for HIVTree. Sequences are shown in the same order in each panel. The vertical lines show the time of infection (dashed), time of treatment start (solid) and the time of sampling (dotted). The color shows the number of RNA sequences subsampled from the original alignment at each sample time. If fewer sequences were available then the number indicated by the color at a given time, all available sequences were used. Sites with greater than 85% missing gaps have been removed from the alignment.

**Fig. S20.**
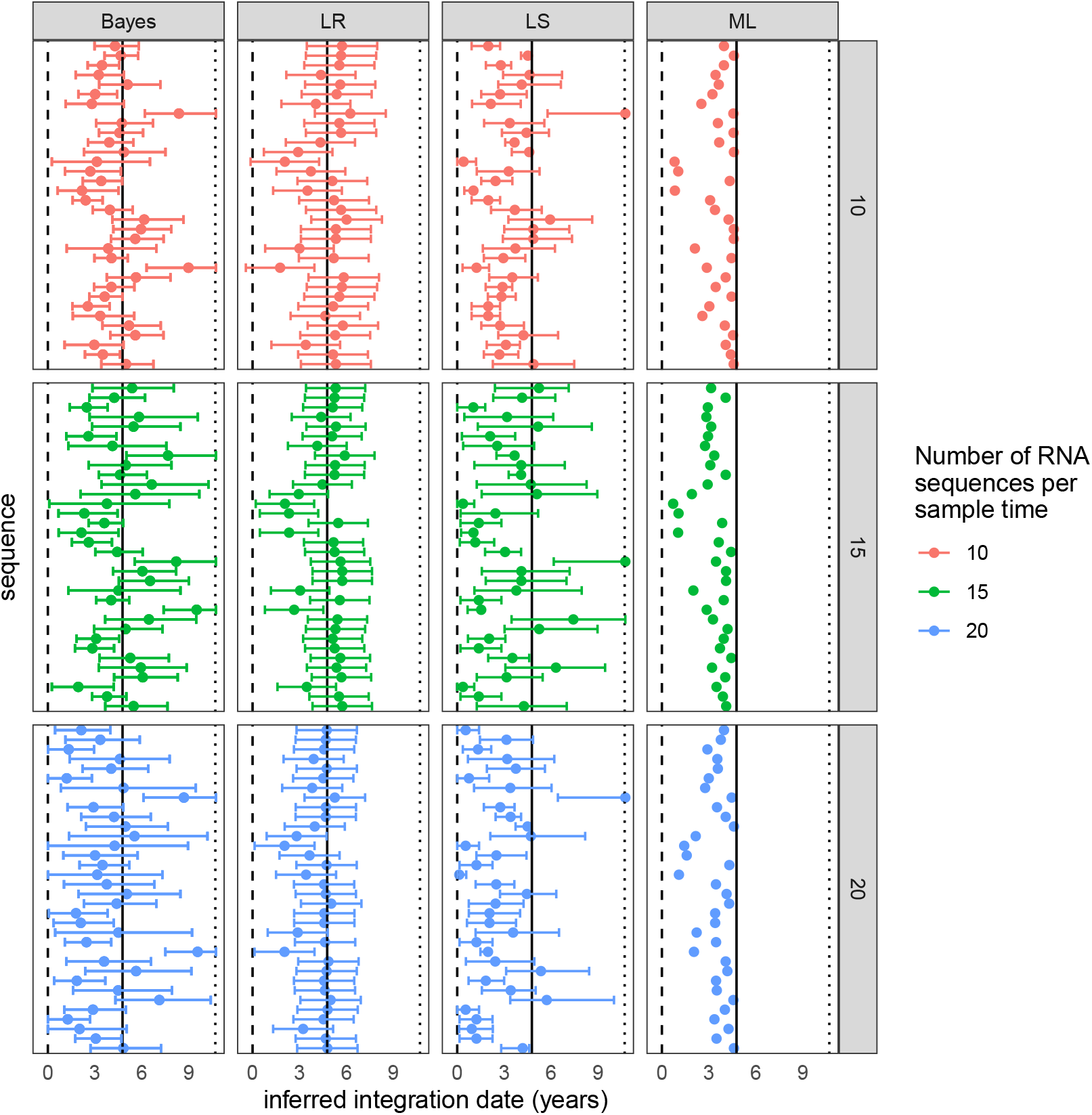
The inferred latent integration dates for Env_4 from patient 257 are shown for each method. 95% confidence intervals are shown for the LR and LS methods, and the 95% credible interval is shown for HIVTree. Sequences are shown in the same order in each panel. The vertical lines show the time of infection (dashed), time of treatment start (solid) and the time of sampling (dotted). The color shows the number of RNA sequences subsampled from the original alignment at each sample time. If fewer sequences were available then the number indicated by the color at a given time, all available sequences were used. Sites with greater than 95% missing gaps have been removed from the alignment.

**Fig. S21.**
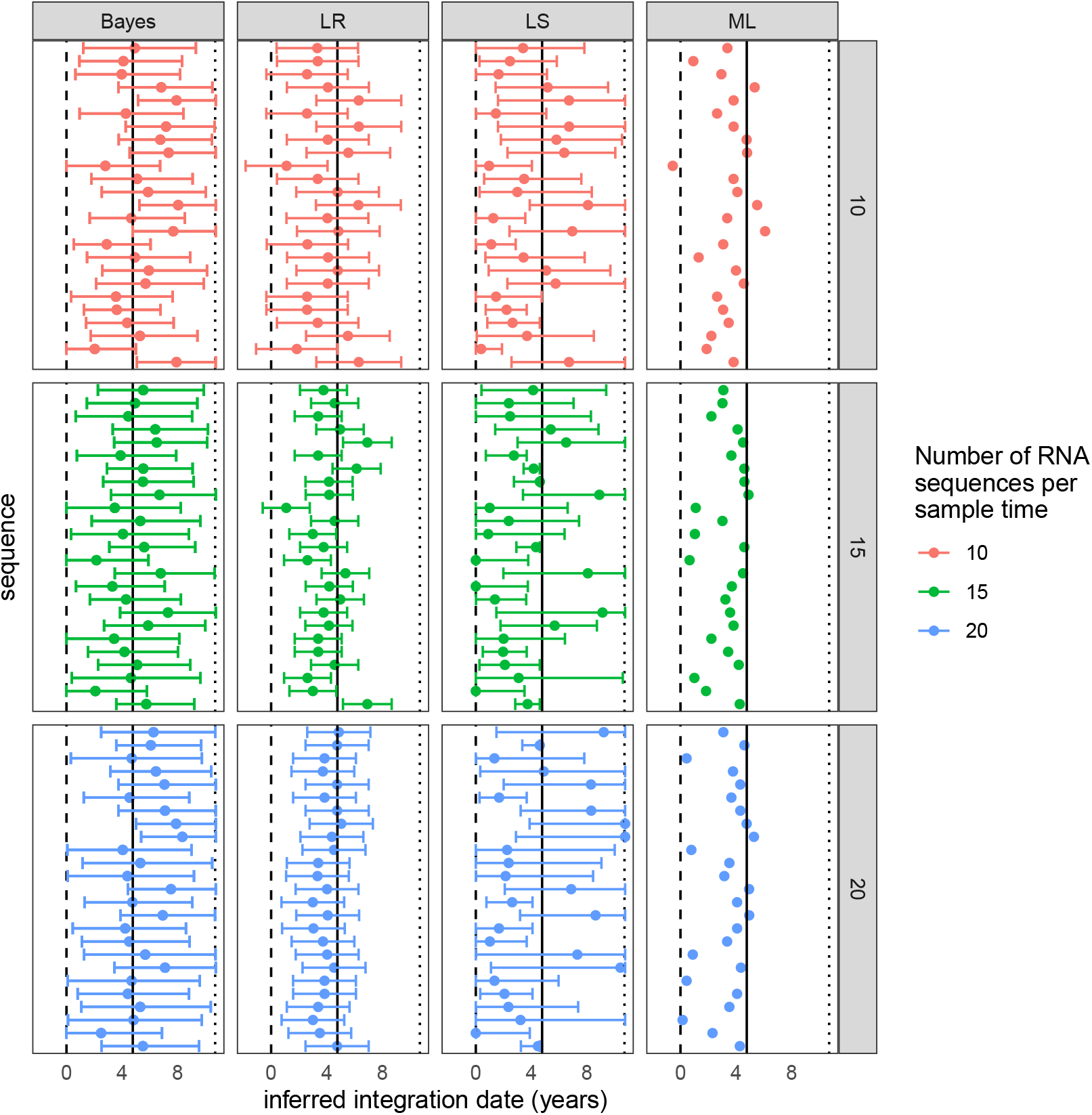
The inferred latent integration dates for GAG_1 from patient 257 are shown for each method. 95% confidence intervals are shown for the LR and LS methods, and the 95% credible interval is shown for HIVTree. Sequences are shown in the same order in each panel. The vertical lines show the time of infection (dashed), time of treatment start (solid) and the time of sampling (dotted). The color shows the number of RNA sequences subsampled from the original alignment at each sample time. If fewer sequences were available then the number indicated by the color at a given time, all available sequences were used. Sites with greater than 75% missing gaps have been removed from the alignment.

**Fig. S22.**
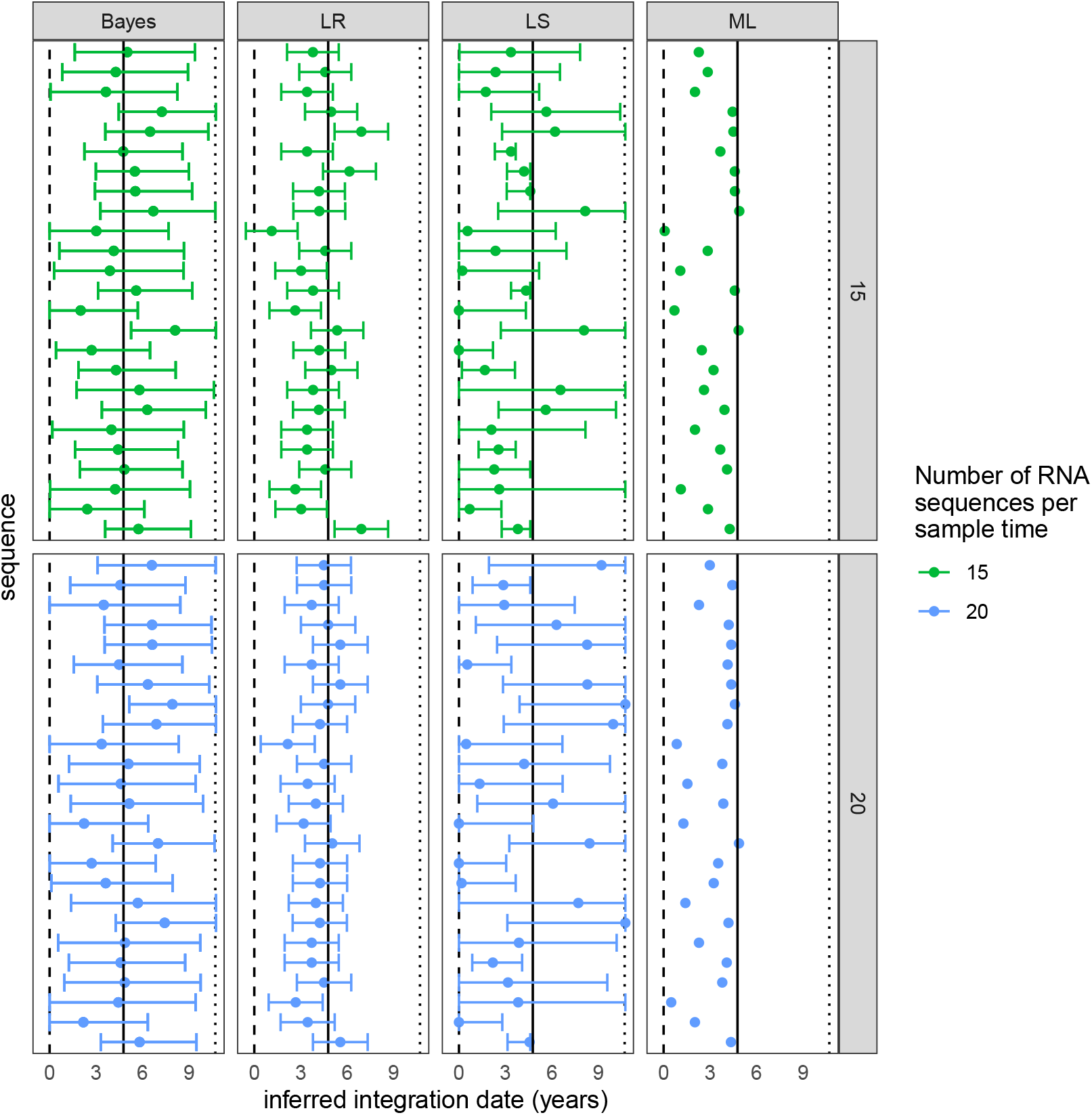
The inferred latent integration dates for GAG_1 from patient 257 are shown for each method. 95% confidence intervals are shown for the LR and LS methods, and the 95% credible interval is shown for HIVTree. Sequences are shown in the same order in each panel. The vertical lines show the time of infection (dashed), time of treatment start (solid) and the time of sampling (dotted). The color shows the number of RNA sequences subsampled from the original alignment at each sample time. If fewer sequences were available then the number indicated by the color at a given time, all available sequences were used. Sites with greater than 95% missing gaps have been removed from the alignment.

**Fig. S23.**
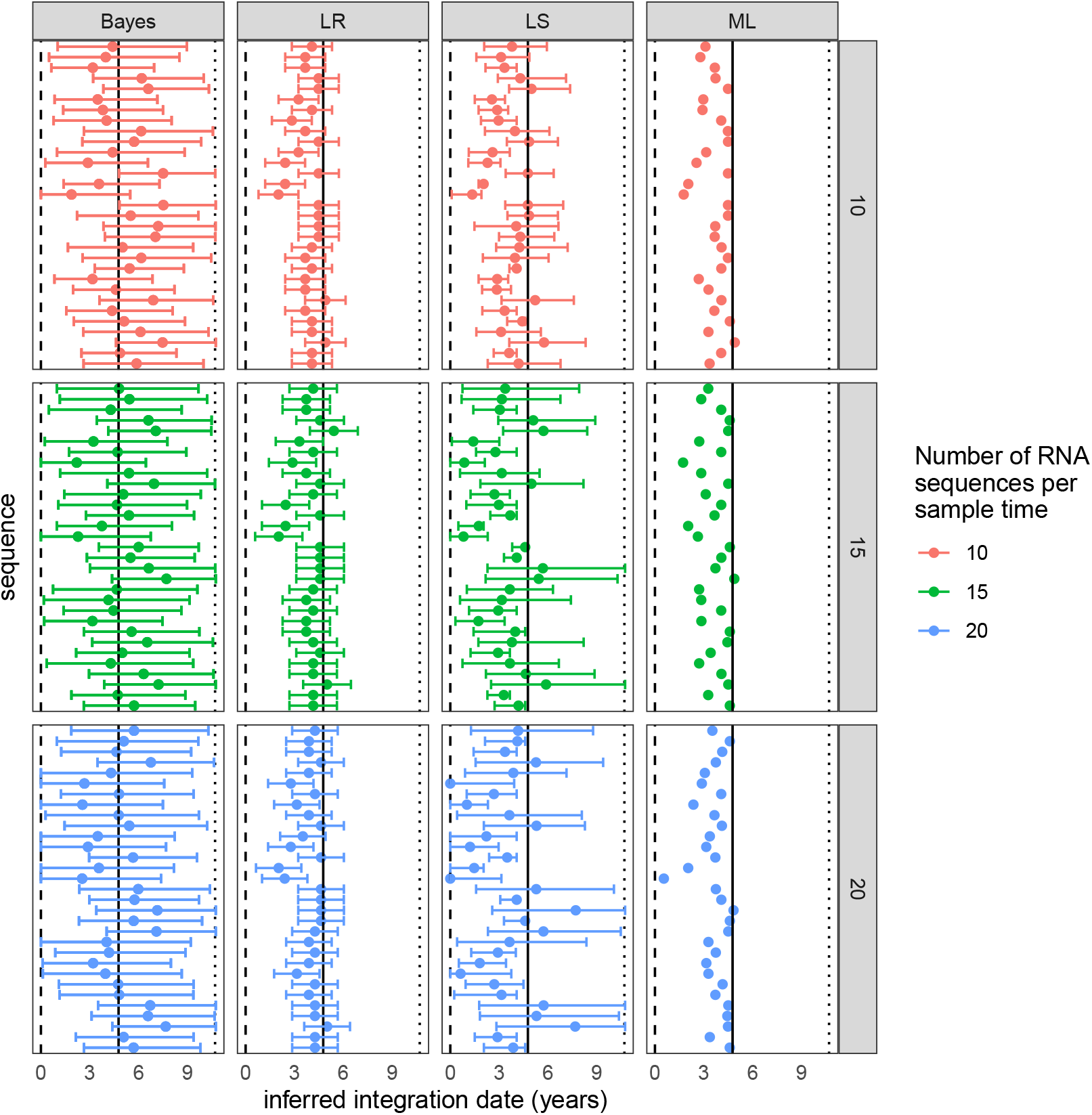
The inferred latent integration dates for NEF_1 from patient 257 are shown for each method. 95% confidence intervals are shown for the LR and LS methods, and the 95% credible interval is shown for HIVTree. Sequences are shown in the same order in each panel. The vertical lines show the time of infection (dashed), time of treatment start (solid) and the time of sampling (dotted). The color shows the number of RNA sequences subsampled from the original alignment at each sample time. If fewer sequences were available then the number indicated by the color at a given time, all available sequences were used. Sites with greater than 75% missing gaps have been removed from the alignment.

**Fig. S24.**
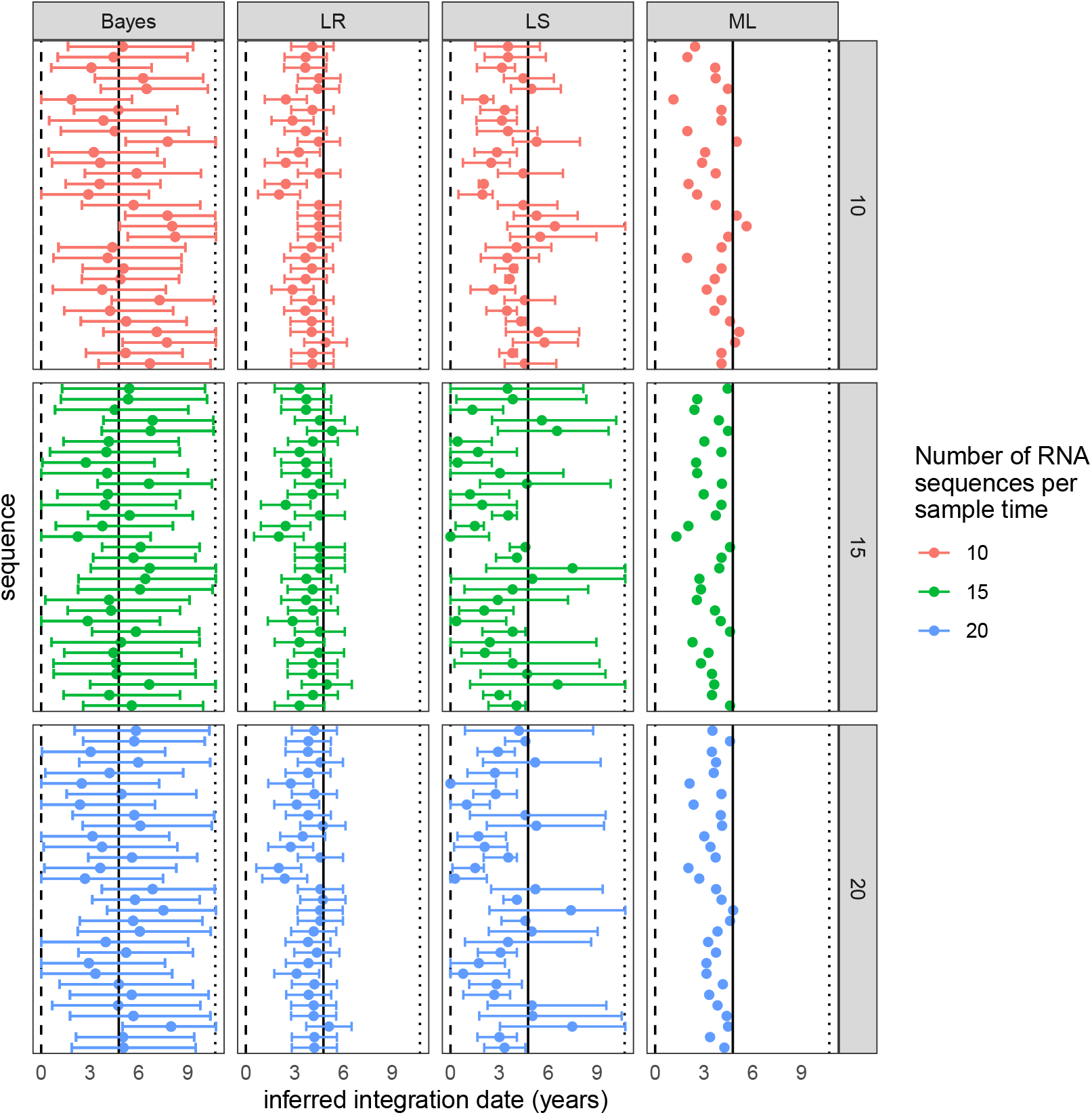
The inferred latent integration dates for NEF_1 from patient 257 are shown for each method. 95% confidence intervals are shown for the LR and LS methods, and the 95% credible interval is shown for HIVTree. Sequences are shown in the same order in each panel. The vertical lines show the time of infection (dashed), time of treatment start (solid) and the time of sampling (dotted). The color shows the number of RNA sequences subsampled from the original alignment at each sample time. If fewer sequences were available then the number indicated by the color at a given time, all available sequences were used. Sites with greater than 95% missing gaps have been removed from the alignment.

**Fig. S25.**
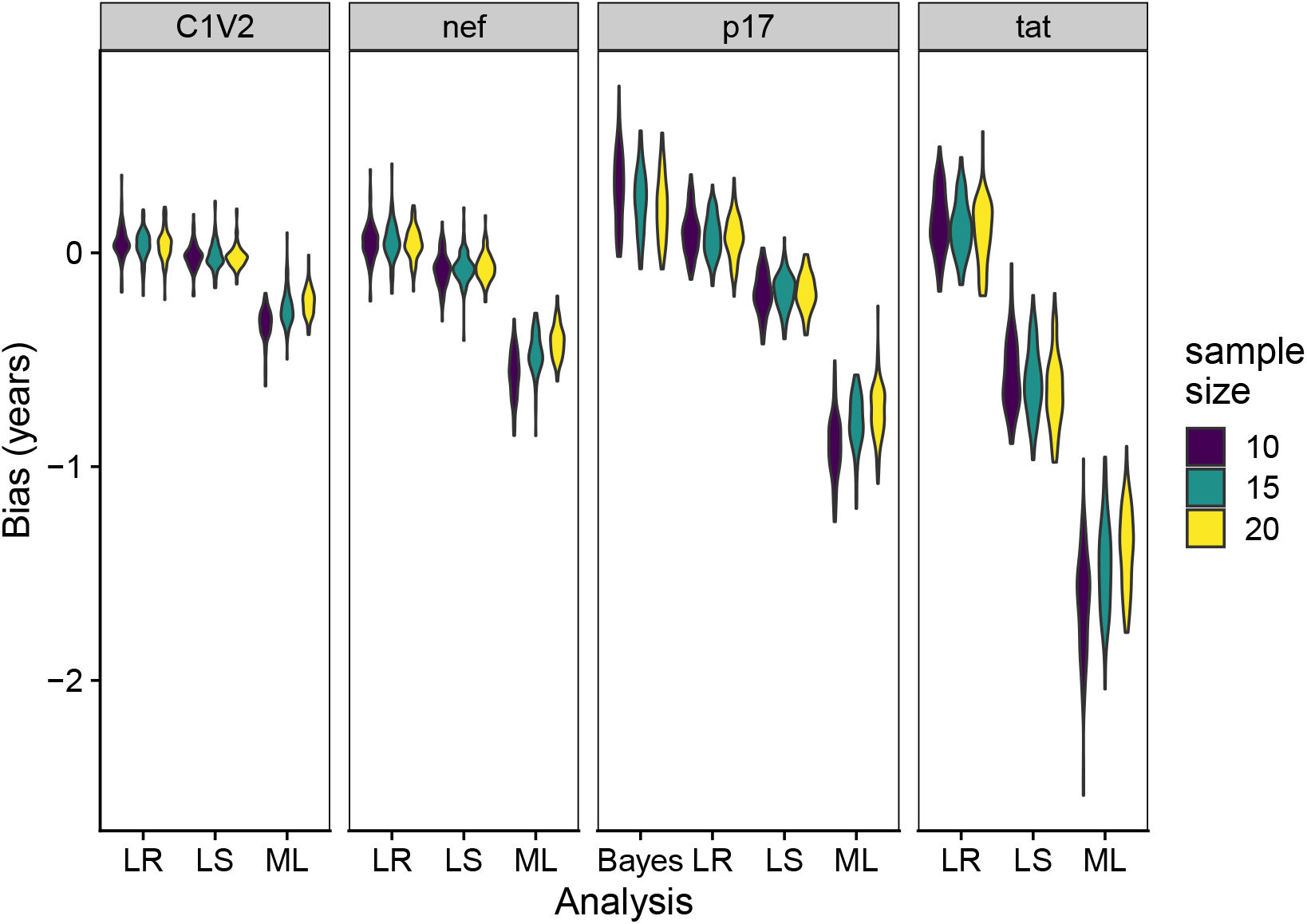
The bias for each simulated region using each of four analysis is shown. Each data point in the violin plot is the average bias of 30 latent times in each of 30 alignments with a fixed topology. There are a total of 100 fixed topolgies for each violin plot. The number of non-latent sequences sampled at each of 10 sampling time points is indicated by the color. While the longest and most quickly evolving gene, *C1V2*, has the lowest bias for all methods and the shorter, more slowly evolving genes have greater bias, there is not a consistent trend in bias by the sample size.

**Fig. S26.**
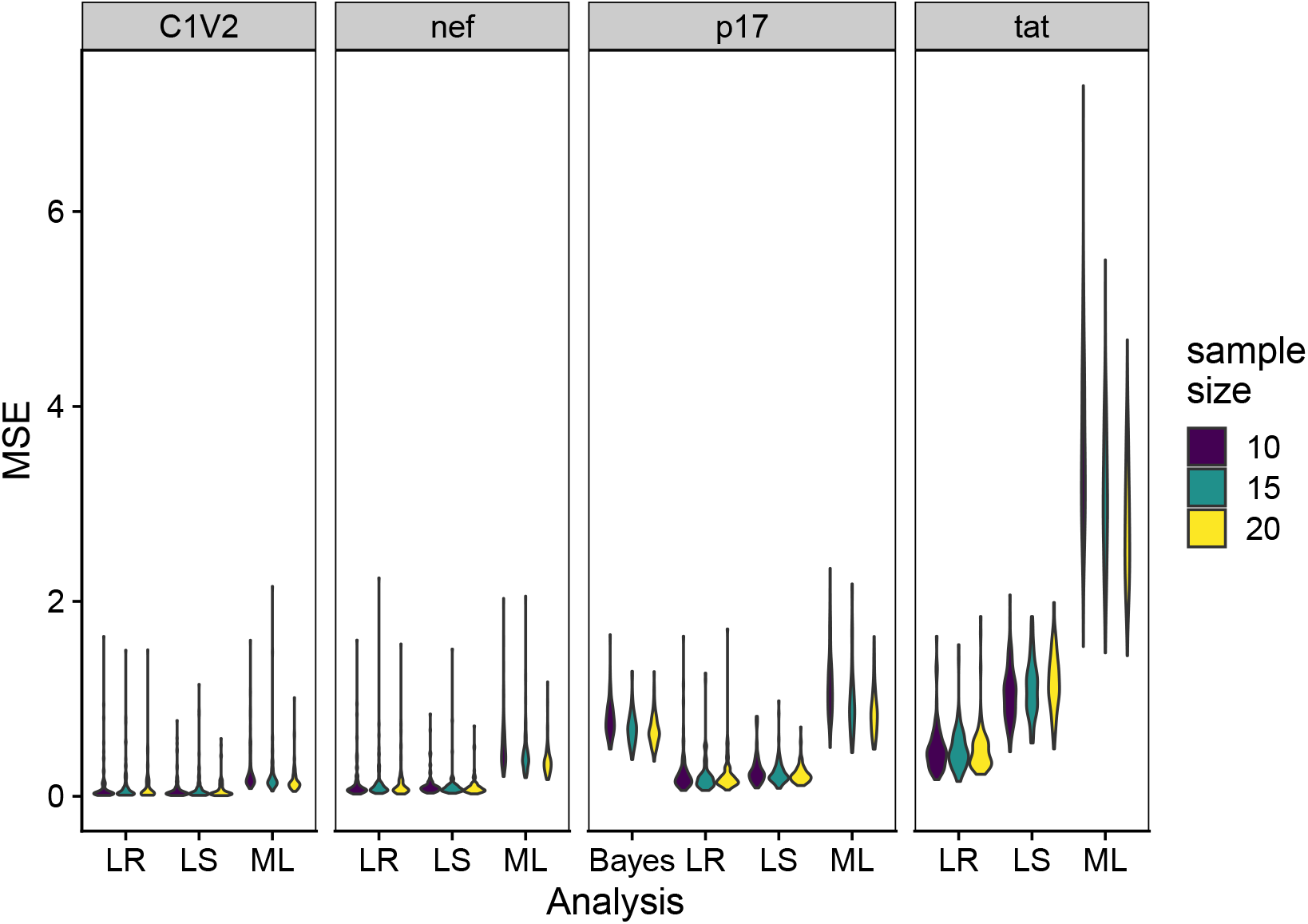
Each data point in the violin plot is the average MSE of 30 latent times in each of 30 alignments with a fixed topology. There are a total of 100 fixed topolgies for each violin plot. The number of non-latent sequences sampled at each of 10 sampling time points is indicated by the color. There is not a consistent trend in MSE by the sample size.

**Fig. S27.**
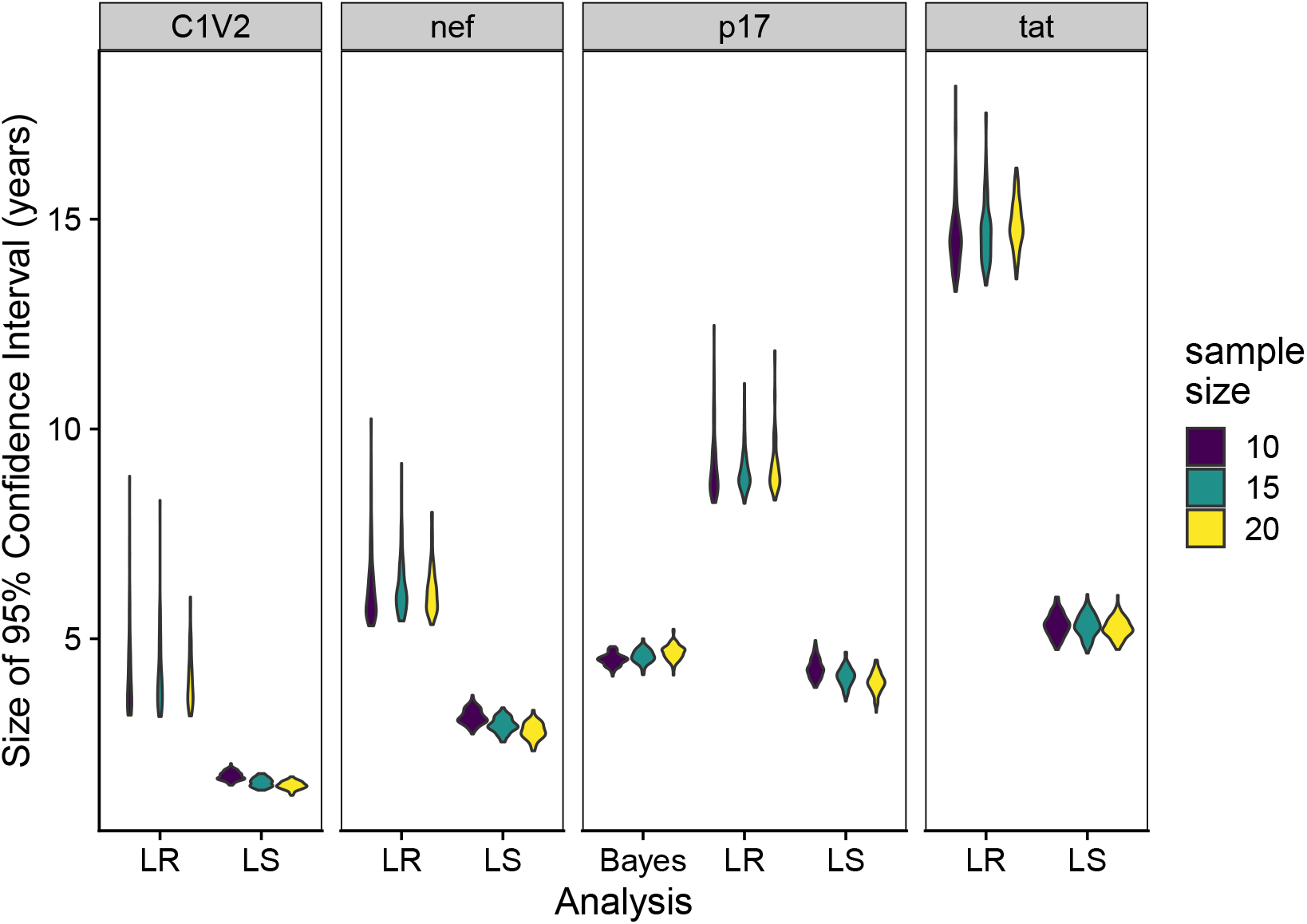
Each data point in the violin plot is the average size of the 95% confidence intervals (or credible sets for the Bayesian method) of 30 latent times in each of 30 alignments with a fixed topology. There are a total of 100 fixed topolgies for each violin plot. The number of non-latent sequences sampled at each of 10 sampling time points is indicated by the color. The longest and most quickly evolving gene, *C1V2*, has smaller confidence intervals for all methods. The sample size does not have a large effect on the size of the confidence intervals.

**Fig. S28.**
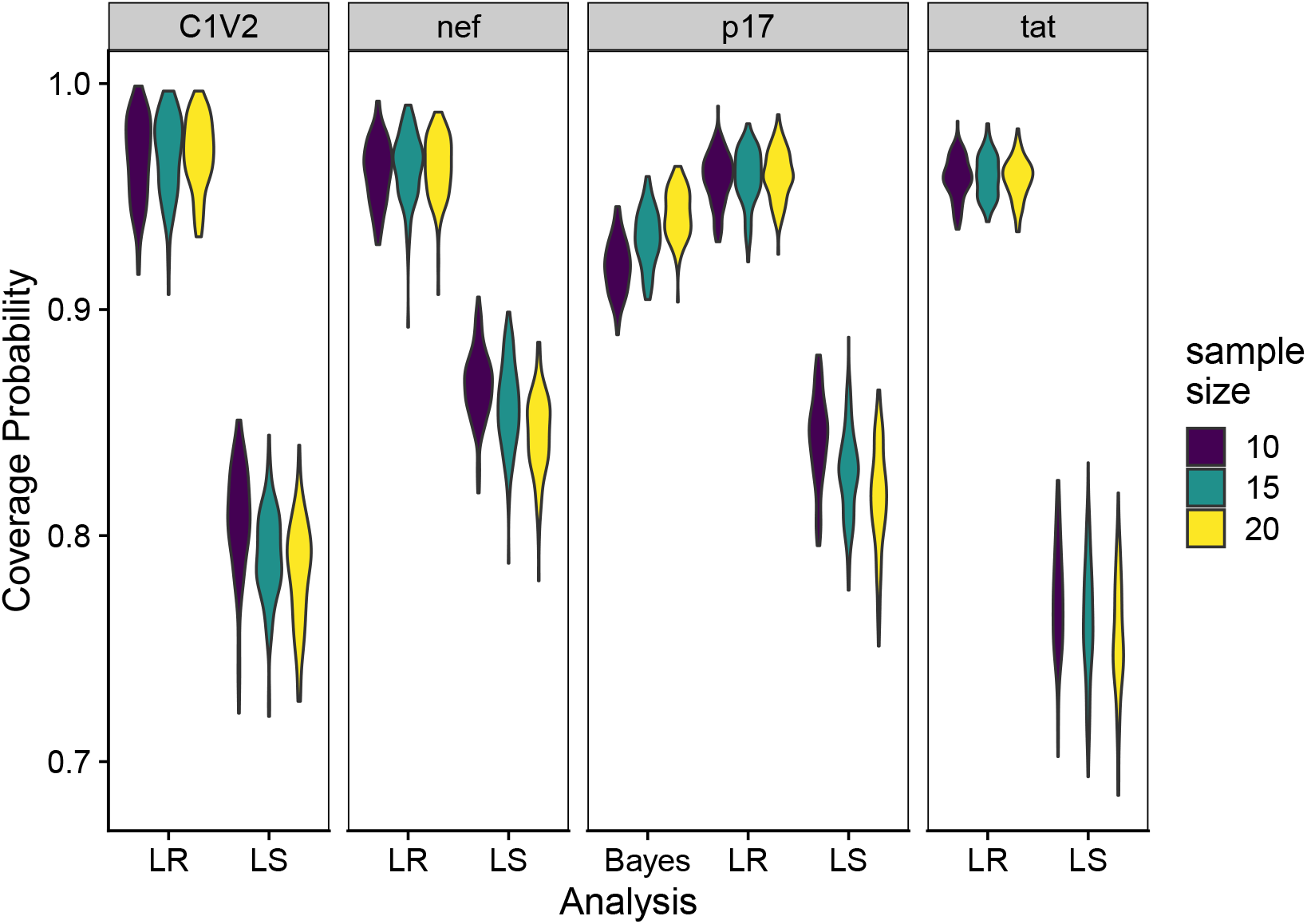
Each data point in the violin plot is the probability the true latent time falls within the 95% confidence intervals (or 95% highest posterior density set) for 30 latent times in each of 30 alignments with a fixed topology. There are a total of 100 fixed topolgies for each violin plot. The number of non-latent sequences sampled at each of 10 sampling time points is indicated by the color. This probability is always 1 for the LR method. For the LS method, the probability decreases when the region is shorter with a lower mutation rate, but does not vary predictably with sample size. The ML method is not shown since it does provide confidence intervals or credible sets.

## References

1. RT Davey, et al., HIV-1 and T cell dynamics after interruption of highly active antiretroviral therapy (HAART) in patients with a history of sustained viral suppression. Proc. Natl. Acad. Sci. 96, 15109–15114 (1999).

2. DD Ho, et al., Rapid turnover of plasma virions and CD4 lymphocytes in HIV-1 infection. Nature 373, 123–126 (1995).

3. X Wei, et al., Viral dynamics in human immunodeficiency virus type 1 infection. Nature 373, 117–122 (1995).

4. JD Siliciano, et al., Long-term follow-up studies confirm the stability of the latent reservoir for HIV-1 in resting CD4 +T cells. Nat. Medicine 9, 727–728 (2003).

5. C Dufour, P Gantner, R Fromentin, N Chomont, The multifaceted nature of HIV latency. J. Clin. Investig. 130, 3381–3390 (2020).

6. RF Siliciano, WC Greene, HIV latency. Cold Spring Harb. Perspectives Medicine 1, a007096 (2011).

7. TW Chun, et al., Early establishment of a pool of latently infected, resting CD4+ T cells during primary HIV-1 infection. Proc. Natl. Acad. Sci. 95, 8869–8873 (1998).

8. JB Whitney, et al., Rapid seeding of the viral reservoir prior to SIV viraemia in rhesus monkeys. Nature 512, 74–77 (2014).

9. C Verhofstede, et al., Drug-resistant variants that evolve during nonsuppressive therapy persist in HIV-1–infected peripheral blood mononuclear cells after long-term highly active antiretroviral therapy. J. Acquir. Immune Defic. Syndr. 35, 473–483 (2004).

10. J Brodin, et al., Establishment and stability of the latent HIV-1 DNA reservoir. Elife 5, e18889 (2016).

11. MR Abrahams, et al., The replication-competent HIV-1 latent reservoir is primarily established near the time of therapy initiation. Sci. Transl. Medicine 11, eaaw5589 (2019).

12. MD Pankau, et al., Dynamics of HIV DNA reservoir seeding in a cohort of superinfected Kenyan women. PLOS Pathog. 16, e1008286 (2020).

13. BR Jones, et al., Phylogenetic approach to recover integration dates of latent HIV sequences within-host. Proc. Natl. Acad. Sci. 115, E8958–E8967 (2018).

14. KM Bruner, et al., Defective proviruses rapidly accumulate during acute HIV-1 infection. Nat. Medicine 22, 1043–1049 (2016).

15. BR Jones, JB Joy, Simulating within host human immunodeficiency virus 1 genome evolution in the persistent reservoir. Virus Evol. 6 (2020).

16. BR Jones, AFY Poon, node.dating: dating ancestors in phylogenetic trees in R. Bioinformatics 33, 932–934 (2017).

17. TH To, M Jung, S Lycett, O Gascuel, Fast dating using least-squares criteria and algorithms. Syst. Biol. 65, 82–97 (2016).

18. T Stadler, Z Yang, Dating phylogenies with sequentially sampled tips. Syst. Biol. 62, 674–688 (2013).

19. A Rambaut, Estimating the rate of molecular evolution: incorporating non-contemporaneous sequences into maximum likelihood phylogenies. Bioinformatics 16, 395–399 (2000).

20. SG Deeks, J Overbaugh, A Phillips, S Buchbinder, HIV infection. Nat. Rev. Dis. Primers 1, 1–22 (2015).

21. AN Phillips, Reduction of HIV concentration during acute infection: Independence from a specific immune response. Science 271, 497–499 (1996).

22. MA Nowak, CRM Bangham, Population dynamics of immune responses to persistent viruses. Science 272, 74–79 (1996).

23. AS Perelson, RM Ribeiro, Modeling the within-host dynamics of HIV infection. BMC Biol. 11, 96 (2013).

24. MA Stafford, et al., Modeling plasma virus concentration during primary HIV infection. J. Theor. Biol. 203, 285–301 (2000).

25. B Rannala, Conceptual issues in Bayesian divergence time estimation. Philos. Transactions Royal Soc. B: Biol. Sci. 371, 20150134 (2016).

26. T Flouri, X Jiao, B Rannala, Z Yang, Species tree inference with BPP using genomic sequences and the multispecies coalescent. Mol. Biol. Evol. 35, 2585–2593 (2018).

27. Z Yang, B Rannala, Bayesian estimation of species divergence times under a molecular clock using multiple fossil calibrations with soft bounds. Mol. Biol. Evol. 23, 212–226 (2006).

28. AG Rodrigo, J Felsenstein, The Evolution of HIV. (The John Hopkins University Press), pp. 233–267 (1999).

29. VN Minin, EW Bloomquist, MA Suchard, Smooth skyride through a rough skyline: Bayesian coalescent-based inference of population dynamics. Mol. Biol. Evol. 25, 1459–1471 (2008).

30. S Tavaré, et al., Some probabilistic and statistical problems in the analysis of DNA sequences. Lect. on mathematics life sciences 17, 57–86 (1986).

31. Z Yang, Maximum-likelihood estimation of phylogeny from DNA sequences when substitution rates differ over sites. Mol. biology evolution 10, 1396–1401 (1993).

32. M Hasegawa, H Kishino, T Yano, Dating of the human-ape splitting by a molecular clock of mitochondrial DNA. J. Mol. Evol. 22, 160–174 (1985).

33. AM Kozlov, D Darriba, T Flouri, B Morel, A Stamatakis, RAxML-NG: a fast, scalable and userfriendly tool for maximum likelihood phylogenetic inference. Bioinformatics 35, 4453–4455 (2019).

34. E Paradis, J Claude, K Strimmer, APE: Analyses of phylogenetics and evolution in R language. Bioinformatics 20, 289–290 (2004).

35. J Moss, M Tveten, kdensity: An R package for kernel density estimation with parametric starts and asymmetric kernels. J. Open Source Softw. 4, 1566 (2019).

36. JM Pavia, Testing goodness-of-fit with the kernel density estimator: GoFKernel. J. Stat. Softw. 66, 1–27 (2015).

## References

1. K Soetaert, T Petzoldt, RW Setzer, Solving differential equations in R: Package deSolve. J. Stat. Softw. 33 (2010).

2. MA Stafford, et al., Modeling plasma virus concentration during primary HIV infection. J. Theor. Biol. 203, 285–301 (2000).

3. TW Chun, et al., Quantification of latent tissue reservoirs and total body viral load in HIV-1 infection. Nature 387, 183–188 (1997).

4. Z Yang, PAML 4: Phylogenetic analysis by maximum likelihood. Mol. Biol. Evol. 24, 1586–1591 (2007).

5. Y Liu, JP McNevin, S Holte, MJ McElrath, JI Mullins, Dynamics of viral evolution and CTL responses in HIV-1 infection. PloS One 6, e15639 (2011).

6. Y Liu, et al., Evolution of human immunodeficiency virus type 1 cytotoxic T-lymphocyte epitopes: fitness-balanced escape. J. Virol. 81, 12179–12188 (2007).

7. Y Liu, et al., Selection on the human immunodeficiency virus type 1 proteome following primary infection. J. Virol. 80, 9519–9529 (2006).

8. AM Kozlov, D Darriba, T Flouri, B Morel, A Stamatakis, RAxML-NG: a fast, scalable and user-friendly tool for maximum likelihood phylogenetic inference. Bioinformatics 35, 4453–4455 (2019).

9. M Hasegawa, H Kishino, T Yano, Dating of the human-ape splitting by a molecular clock of mitochondrial DNA. J. Mol. Evol. 22, 160–174 (1985).

10. Z Yang, Maximum-likelihood estimation of phylogeny from DNA sequences when substitution rates differ over sites. Mol. biology evolution 10, 1396–1401 (1993).

11. T Stadler, Z Yang, Dating phylogenies with sequentially sampled tips. Syst. Biol. 62, 674–688 (2013).

12. Z Yang, B Rannala, Bayesian estimation of species divergence times under a molecular clock using multiple fossil calibrations with soft bounds. Mol. Biol. Evol. 23, 212–226 (2006).

14. J Rozewicki, S Li, KM Amada, DM Standley, K Katoh, MAFFT-DASH: integrated protein sequence and structural alignment. Nucleic Acids Res. 47, W5–W10 (2019).

15. MR Abrahams, et al., The replication-competent HIV-1 latent reservoir is primarily established near the time of therapy initiation. Sci. Transl. Medicine 11, eaaw5589 (2019).

16. R Luo, MJ Piovoso, J Martinez-Picado, R Zurakowski, HIV model parameter estimates from interruption trial data including drug efficacy and reservoir dynamics. PLoS ONE 7, e40198 (2012).

17. AL Hill, DIS Rosenbloom, F Fu, MA Nowak, RF Siliciano, Predicting the outcomes of treatment to eradicate the latent reservoir for HIV-1. Proc. Natl. Acad. Sci. 111, 13475–13480 (2014).

18. KM Bruner, et al., Defective proviruses rapidly accumulate during acute HIV-1 infection. Nat. Medicine 22, 1043–1049 (2016).

19. MJ Peluso, et al., Differential decay of intact and defective proviral DNA in HIV-1–infected individuals on suppressive antiretroviral therapy. JCI Insight 5, e132997 (2020).

